# Genetic Impairment of Succinate Metabolism Disrupts Bioenergetic Sensing in Adrenal Neuroendocrine Cancer

**DOI:** 10.1101/2022.01.09.475410

**Authors:** Priyanka Gupta, Keehn Strange, Rahul Telange, Ailan Guo, Heather Hatch, Amin Sobh, Jonathan Elie, Angela M. Carter, John Totenhagen, Chunfeng Tan, Yogesh A. Sonawane, Jiri Neuzil, Amarnath Natarajan, Ashley J. Ovens, Jonathan S. Oakhill, Thorsten Wiederhold, Karel Pacak, Hans K. Ghayee, Laurent Meijer, Sushanth Reddy, James A. Bibb

## Abstract

Metabolic dysfunction mutations can impair energy sensing and cause cancer. Loss of function of mitochondrial TCA cycle enzyme, succinate dehydrogenase B (SDHB) results in various forms of cancer typified by pheochromocytoma (PC). Here we delineate a signaling cascade where the loss of *SDHB* induces the Warburg effect in PC tumors, triggers dysregulation of Ca^2+^ homeostasis, and aberrantly activates calpain and the protein kinase Cdk5, through conversion of its cofactor from p35 to p25. Consequently, aberrant Cdk5 initiates a cascade of phospho-signaling where GSK3 inhibition inactivates energy sensing by AMP-kinase through dephosphorylation of the AMP-kinase γ subunit, PRKAG2. Overexpression of p25-GFP in mouse adrenal chromaffin cells also elicits this phosphorylation signaling and causes PC tumor formation. A novel Cdk5 inhibitor, MRT3-007, reversed this phospho-cascade, invoking an anti-Warburg effect, cell cycle arrest, and senescence-like phenotype. This therapeutic approach halted tumor progression *in vivo*. Thus, we reveal an important novel mechanistic feature of metabolic sensing and demonstrate that its dysregulation underlies tumor progression in PC and likely other cancers.

**Highlights:**

- Loss of SDHB function in pheochromocytoma causes Ca^2+^ dysregulation, calpain activation, and aberrant activation of the protein kinase Cdk5.
- Hyperactive Cdk5 deregulates a GSK3/PRKAG2/AMPKα signaling cascade.
- p25 overexpression and consequent aberrant Cdk5 activity in chromaffin cells causes pheochromocytoma.
- Inhibition of Cdk5 activates the PRKAG2/AMPK/p53 signaling to rescue cell senescence and block PC tumor progression.

**Graphical Abstract:** 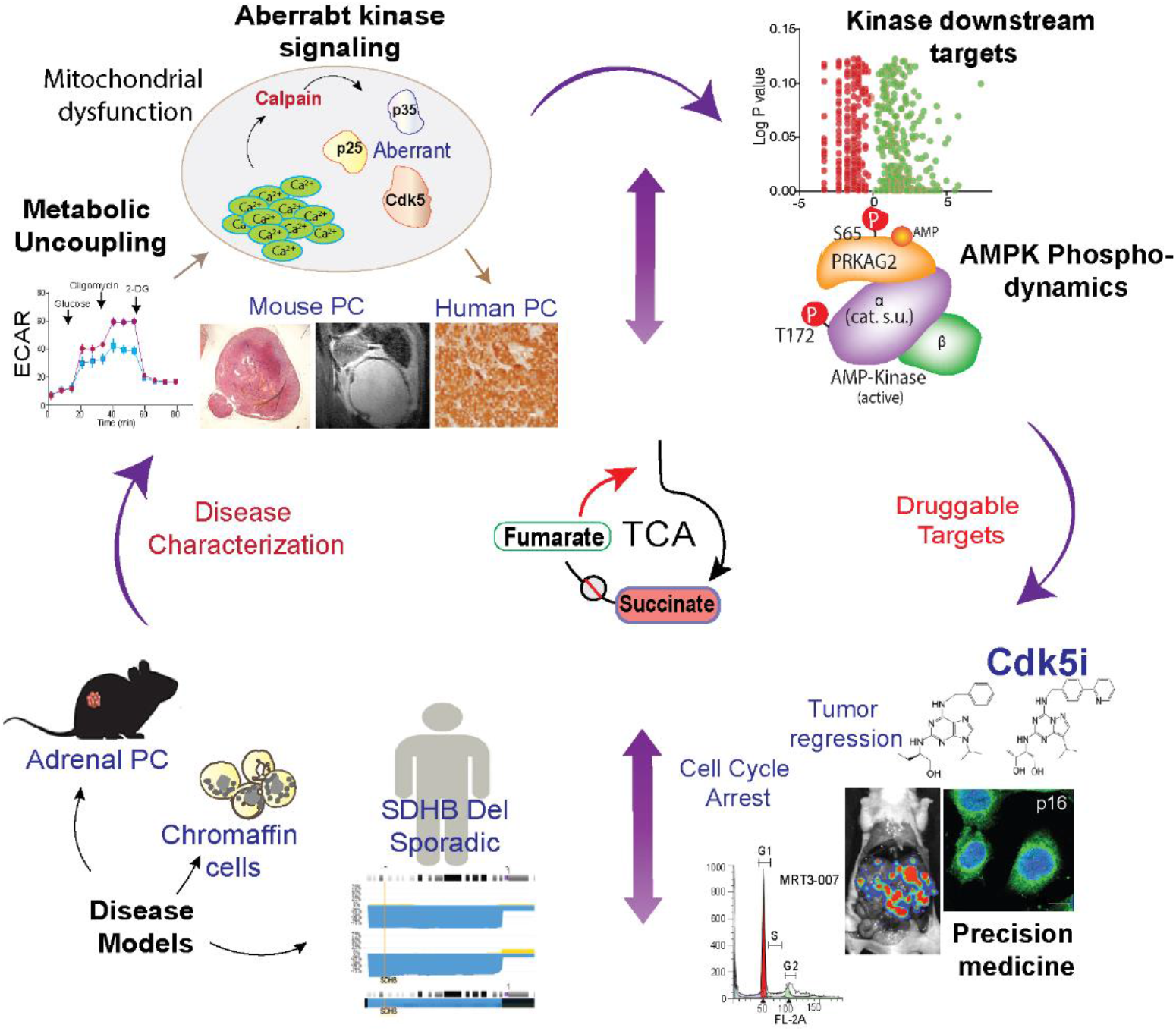

## INTRODUCTION

The tricarboxylic acid (TCA) cycle is the second component of cellular respiration following glycolysis, and the centerpiece of cell metabolism. Significant alterations in TCA cycle components cause a spectrum of metabolic disorders (Krebs and Johnson, 1980). These alterations may be compatible with embryonic and postnatal development but can result in organ-system dysfunction or context-specific pathology (Jeanmonod et al., 2020; Stockholm et al., 2005). Multiple enzymes of the TCA cycle such as fumarate hydratase *(FH),* isocitrate dehydrogenase *(IDH) and* succinate dehydrogenase *(SDH)* are altered in numerous sporadic or hereditary forms of cancer, causing production of oncometabolites. Pheochromocytoma (PC) and paraganglioma (PG) are archetypical metabolic diseases in which TCA malfunction results in tumors arising from chromaffin cells of the adrenal medulla or sympathetic and parasympathetic ganglia (Dahia, 2014; Erez et al., 2011). Mutations in succinate dehydrogenase complex genes (*SDHx*) associate with high incidence of the disease. Moreover, high rates of *SDHx*-related abnormalities correlate with increased risk of metastatic PC/PG in patients who developed primary tumors in childhood or adolescence (King et al., 2011).

The *SDHx* complex is a hetero-tetrameric protein composed of four subunits- SDHA, SDHB, SDHC, and SDHD that functions in both the TCA cycle, and electron transport chain (ETC) catalyzing the oxidation of succinate to fumarate, and converting ubiquinone to ubiquinol, respectively (Bardella et al., 2011). Patients harboring mutations in *SDHB* have a lifetime cancer risk of 76%, with 50% penetrance by the age of 35 (Neumann et al., 2004). A wide array of intragenic mutations are reported in *SDHx* complex genes including frameshifts, splicing defects, and recently reported single/multiple exon deletions. Approximately 60% of *SDHB*-related adrenal or extra-adrenal PC present allelic imbalances or somatic deletions of wild-type (WT) alleles (Gimenez-Roqueplo et al., 2003; Gimenez-Roqueplo et al., 2002). A significant focal deletion in the *SDHB* locus of chromosome 1 in 91 out of 93 PC/PG patients was identified in a publicly available human PG/PG dataset (NCIC3326) (https://progenetix.org/progenetix-cohorts/TCGA/). These defects in *SDHB* exhibit complicated phenotypes that affect gene expression and enzymatic function to varying degrees (Huang et al., 2018; Solis et al., 2009).

Malfunction due to loss of *SDHx* genes *causes* abnormal accumulation of succinate, thereby activating angiogenic/hypoxia-responsive genes such as *EPAS1, HIF1α, and VEGF*. These effectors further promote metabolic re-programming and tumor progression (Anderson et al., 2018). Other malignancies affected by *SDHB* loss of function include renal carcinoma, gastrointestinal stromal tumors, pancreatic neuroendocrine tumors, pituitary adenoma, and pulmonary chondroma (Eijkelenkamp et al., 2020). For all these diseases, silencing *SDHB* impairs mitochondrial respiration and causes metabolic shift in favor of aerobic glycolysis to meet the high energetic and biosynthetic demands of tumor cells, a process known as the Warburg effect (Favier et al., 2009; Kim and Dang, 2006)

Impaired TCA cycle function due to *SDHB* loss not only perturbs cellular bioenergetics but also elicits tumor-promoting signaling mediated by Ca^2+^ or reactive oxygen species (ROS) released by mitochondria (Hadrava Vanova et al., 2020; Jana et al., 2019). Altered metabolism or periodic activation of metabolic enzymes maintains a constant supply of ATP and promulgates protein synthesis required for G1/S transition, chromosomal segregation, and activation of cyclin-CDKs complexes (Icard et al., 2019). Thus, TCA impairment results in override of cell cycle checkpoint regulators. Cells may become poorly differentiated, and malignant cell cycle progression can ensue.

While the loss of *SDHB* characterizes precarious forms of PC and other diseases (Jochmanova and Pacak, 2016), the underlying signal transduction mechanisms by which metabolism is reprogrammed and connected to the loss of cell cycle control is not yet fully understood. As a result, it has been challenging to model these diseases or demonstrate targets for therapeutic development. Consequently, little progress has been made in improving the outcomes for patients with these metabolic errors. Here we investigated the overall impact of metabolic dysfunction caused by loss of SDHB on bioenergetics, Ca^2+^ dynamics, neuronal kinase signaling, and tumor progression. These studies provide a new understanding of the determinants of kinase signaling which controls proliferation and senescence-like features, subsequently revealing drug targets for the treatment of PC and other *SDHB* mutation-derived disorders.

## RESULTS

### *SDHB* loss alters cellular metabolism and dysregulates Cdk5 signaling

Genetic alterations in *SDHx* predispose to several metabolic diseases for which the underlying mechanisms are incompletely understood. Heterozygous *SDHB* deletions/mutations are often associated with reduced mRNA expression, or truncated protein (Cascon et al., 2006; Huang et al., 2018; Yang et al., 2012), which may result in impaired but not necessarily complete loss of SDH activity. Analysis of a PCPG dataset revealed genetic alterations within *SDHx* complex in 54% of patients (n=161). The highest portion of 39% occur in *SDHB*/SDHD where majority samples exhibit reduced *SDHB* expression levels (Figure 1A-B). Moreover, approximately 60% of PC patients with shallow deletions in *SDHB* exhibit corresponding reduction in *SDHB* mRNA relative to individuals with unaltered copy of genes (Figure1C). A significant decrease in *SDHB* transcripts in PC patients compared to normal adrenals indicates *SDHB* haploinsuffciency and thus sensitivity to copy number alterations (Figure 1D). Heterozygous deletions/mutations in *SDHB* have greater propensities to induce tumor formation and metastasis (Baysal and Maher, 2015; Buffet et al., 2012). Given these considerations, we conducted CRISPR-Cas9 targeted gene deletion of *SHDB* from progenitor human PC tumor-derived cell line, hPheo1. (Ghayee et al., 2013), and selected a partial knockout (KO) clone expressing 20% of basal *SDHB* levels as an accurate cellular model of the human disease (Figure 1E).

**Figure 1.**
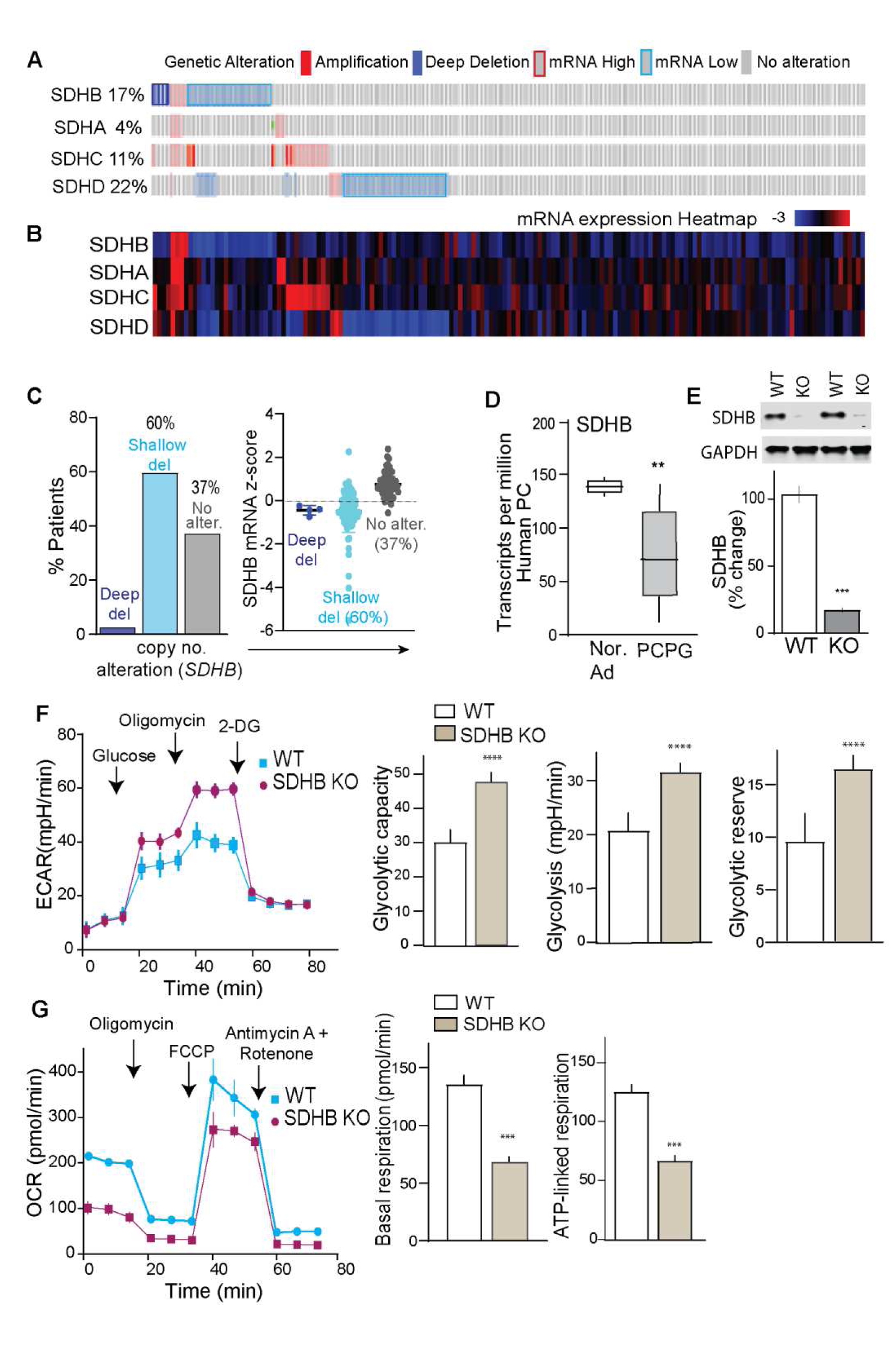
Alterations of *SDHx* genes in pheochromocytoma. (A) Oncoprint depicts alterations in *SDHx* complex genes (SDHB, SDHA, SDHC, SDHD) in 161 PC patient samples. Genetic alterations indicated by color codes. (B) Expression heatmap of *SDHx* complex genes (superimposed on the genetic alterations of the respective samples). Data quarried from cBioportal PCPG PanCancer atlas (http://www.cbioportal.org). (C) Bar plots presenting copy number alteration and mRNA expression of SDHB in PC patient dataset quarried from cBioportal (del., deletions; alter., alterations). (D) Box plots of SDHB gene expression in PC/PG tumors (n=179) vs. normal adrenal (n=3). (E) Exemplary immunoblot showing SDHB protein expression in WT vs*. SDHB* KO hPheo1 cell lysates. (F) Glycolytic profile of WT vs*. SDHB* KO hPheo1 cells indicated as extracellular acidification rate (ECAR) after sequential injections of glucose (10 mM), oligomycin (1µM), and 2-DG (100 mM). Quantitative ECAR analysis indicated as glycolytic capacity, glycolysis and glycolytic reserve. (G) Oxygen consumption rate (OCR) measured in WT and *SDHB* KO cells after sequential addition of Oligomycin (1.5 µM), FCCP (1 µM), and Antimycin + Rotenone (0.5 µM each), respectively. Bar plots are also shown for OCR analysis as basal and ATP-linked respiration, n=3, 10 replicates per group. Values are means ± S.E.M, ***p<0.001, ****p<0.0001 compared by Student’s *t*-test.

Altered metabolism in cancer cells is a common feature where TCA cycle impairment in tumors harboring mutations in *SDHB* or other key components of aerobic respiration contribute to the Warburg effect (Vander Heiden et al., 2009). In agreement, *SDHB* KO dramatically altered the cellular bioenergetics profile. Sequential metabolic perturbation with glucose, oligomycin, and 2-deoxy-D-glucose (2-DG) enhanced the extracellular acidification rate (ECAR), which corresponded to metabolic shifts toward increased glycolysis, glycolytic capacity, and glycolytic reserve in *SDHB* KO cells (Figure 1F). These effects recapitulate the metabolic phenotype that characterizes *SDHB* mutant human PC (Favier et al., 2009; Jochmanova and Pacak, 2016). Concomitant comparison of mitochondrial function indicated elevated oxygen consumption rate (OCR) and higher basal and ATP-linked respiration in WT hPheo1, suggesting decreased efficiency of mitochondrial function due to loss of *SDHB* (Figure 1G). Nevertheless, when challenged with high energy demand via disruption of mitochondrial membrane potential with carbonyl cyanide-4 (trifluoromethoxy) phenylhydrazone (FCCP), a significant spike in maximum respiration was induced in both cell types, suggesting glycolytic preference but not complete shut- off of mitochondrial function in *SDHB* KO PC cells.

Loss of SDHx function not only alters metabolism but also may dysregulates Ca^2+^ homeostasis in neurons (Nasr et al., 2003; Ranganayaki et al., 2021). To evaluate the effect of *SDHB* loss on Ca^2+^ homeostasis in PC, single cell confocal imaging was performed to compare intracellular [Ca^2+^]_i_ in WT vs *SDHB* KO cells using a cell permeable Ca^2+^ indicator. In WT cells, a rapid time-dependent recovery to the ionomycin-induced spike in [Ca^2+^]_i_ occurred, where [Ca^2+^]_i_ surge returning to basal levels within 5 min. In contrast, this response was absent in *SDHB* KO cells where increased levels of [Ca^2+^]_i_ were sustained without significant reduction over the same period of analysis (Figure 2A).

**Figure 2.**
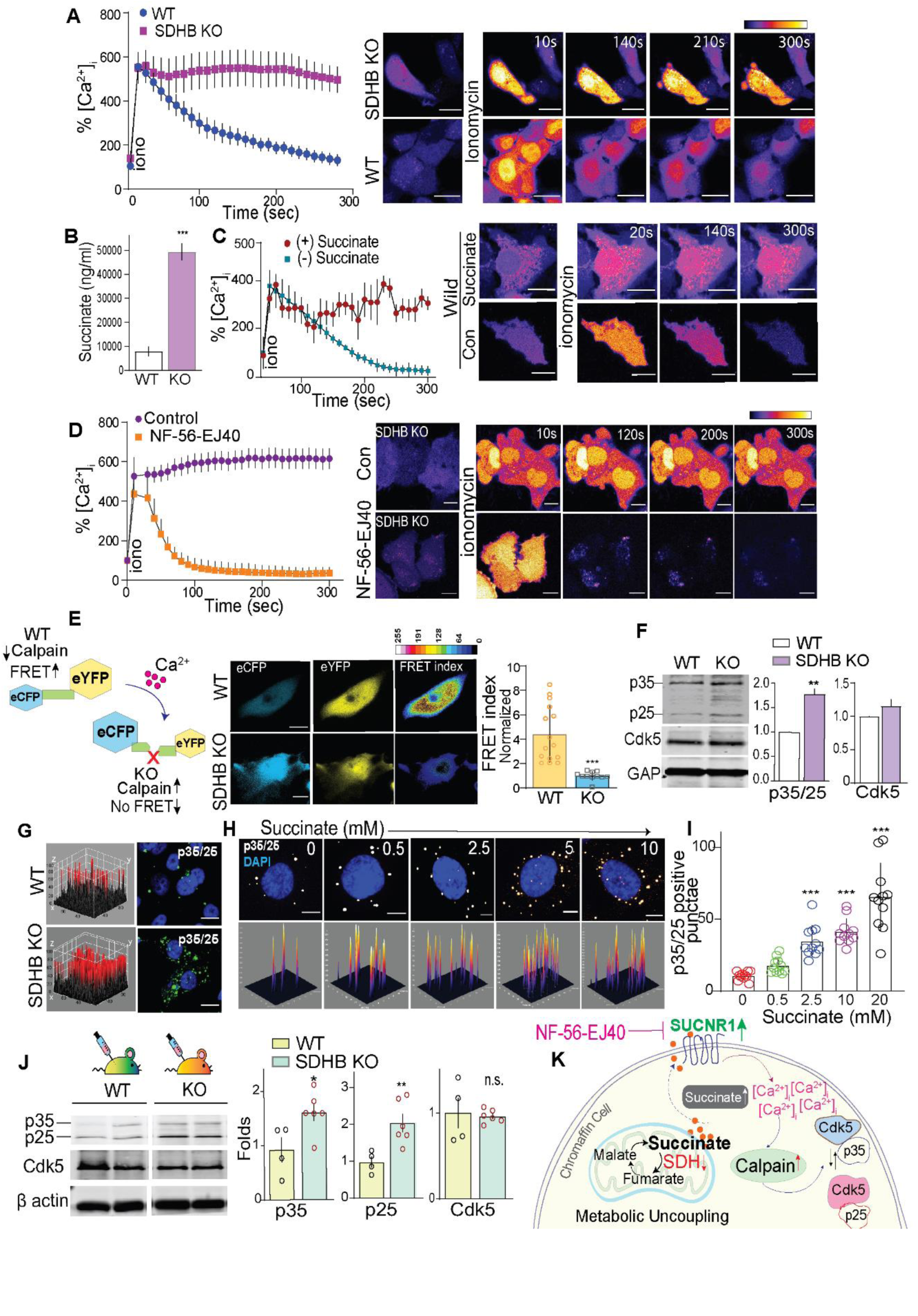
SDHB loss activates a succinate-Ca^2+^-calpain-Cdk5 cascade. (A) [Ca^2+^]_i_ activity reported by time lapse live-cell imaging of PC cells (WT vs*. SDHB* KO) loaded with Fluo-4 AM, images acquired pre- or post-stimulation with ionomycin (10 µM). Representative pseudo-colored images are shown with time-course of fluorescence intensity quantification as % [Ca^2+^]_i_. (B) LC-MS quantitation of succinate in WT and KO cell extracts. (C) Quantitation of fluorescence intensity and photomicrographs showing effects of ionomycin on %[Ca^2+^]_i_ in WT hPheo1 cells treated with succinate (2 mM) or controls. (D) Time lapse measurement of [Ca^2+^]_i_ in *SDHB* KO cells pretreated with SUCNR1 antagonist, NF-56-EJ40, with representative images. (E) Schematic workflow of calpain sensor, and representative pseudocolor FRET map and normalized FRET index of wild type and *SDHB* KO hPheo1 cells. (F) Immunoblots of Cdk5, p25/p35 in WT and *SDHB* KO cells with quantitation. (G) 3D surface plot of immunostained cells comparing relative levels of p35/25 between WT and KO cells. (H-I) Representative confocal 3D photomicrographs and quantitation showing p35/25 expression in WT hPheo1 treated with increasing concentration of succinate as indicated. (J) Quantitative immunoblotting of Cdk5/p35/p25 levels in tumor lysates derived from WT (n=4) and KO (n=6) xenografts. Data are means ± SEM, *p < 0.05, **p < 0.01, ***p<0.001; n.s., non-significant compared by *t*-test and or one-way ANOVA. (K) Schematic illustrating mechanism of action mediated via succinate accumulation and concomitant disruption of Ca^2+^ / Cdk5 homeostasis in chromaffin cells.

In addition to causing an imbalance in [Ca^2+^]_i_ dynamics, *SDHB* loss is commonly thought to trigger build up in the TCA metabolite, succinate. Indeed, succinate concentrations were significantly higher in *SDHB* KO cells compared to controls (Fig. 2B). We hypothesized that this increase in basal intracellular succinate could contribute to the loss in [Ca^2+^]_i_ dynamics that we observed in KO cells. To test this, WT PC cells were treated with cell permeable dimethyl succinate (DMS). Similar to the effect of *SDHB* KO, the addition of exogenous succinate disrupted ionomycin-induced [Ca^2+^]_i_ homoeostasis recovery in parent cells (Figure 2C).

Succinate may alter [Ca^2+^]_i_ homeostasis through autocrine signaling. Specifically, excessive succinate freely shuttles between the mitochondria, cytosol, and across the cell membrane to stimulate the SUCNR1 receptor in *SDHx* related PC tumors (Matlac et al., 2021). Interestingly, hPheo1 cells expressed appreciable levels of SUCNR1 and *SDHB* KO caused a 1.75-fold increase in SUCNR1 expression and significantly elevate intracellular succinate levels (Figure S1A). To test if *SDHB* loss and consequent succinate accumulation triggers SUCNR1 mediated disruption of [Ca^2+^]_i_ homeostasis, KO cells were treated with SUCNR1 antagonist (NF- 56-EJ40). [Ca^2+^]_i_ homeostasis was impaired as was observed earlier in KO cells. However pretreatment with NF-56-EJ40, rescued the WT phenotype for [Ca^2+^]_i_ recovery in *SDHB* KO cells (Figure 2D and S1B). These data indicate that *SDHB* loss caused an increase in succinate which then destabilized [Ca^2+^]_i_ management through the constitutive activation of SUCNR1 receptors.

The Ca^2+^-dependent protease calpain is an important downstream effector activated by loss of Ca^2+^ homeostasis (Crespo-Biel et al., 2007; Nasr et al., 2003; Pang et al., 2003). To determine if *SDHB* KO activated calpain, a FRET (Fluorescence Resonance Energy Transfer) probe harboring a calpain-specific substrate was used (Stockholm et al., 2005). *SDHB* KO triggered an increase in intracellular calpain activity as indicated by significantly higher FRET index in parent hPheo1 vs. *SDHB* KO cells expressing the calpain reporter (Figure 2E and S2A). Consistent with these data, an increase in the 145 kDa breakdown product of spectrin confirmed elevated ubiquitous calpain activity in *SDHB* deficient cells (Figure S3A) (Rajgopal and Vemuri, 2002).

We have shown that Cdk5 is expressed in human PC tumors (Carter et al., 2020). Furthermore, the Cdk5 activating cofactor p35 is an important substrate of calpain that is cleaved to p25 in response to loss of [Ca^2+^]_i_ homeostasis such as that which occurs during neuronal excitotoxicity. The resulting Cdk5/p25 holoenzyme engenders aberrant activity, which can cause neuronal death and may drive neuroendocrine cell proliferation (Barros-Minones et al., 2013; Carter et al., 2020; Crespo-Biel et al., 2009). Quantitative immunoblotting revealed that *SDHB* KO caused a cumulative increase of both p35 and p25 levels compared to WT cells (Fig. 2F). Concomitant immunofluorescent quantitation showed a 22.1 ± 2.4 pixel intensity increase in p35/p25 signals in KO vs. parent cell lines (p=0.002) (Figure 2F-G). Moreover, exogenous succinate induced a dose-dependent increase in p35/p25 levels, with a simultaneous increase in spectrin cleavage in parent cells (Figure 2H-I and S3B).

The increased expression of p35 and p25 generation in response to *SDHB* KO detected in cultured cells was also observed *in vivo* when WT or *SDHB* KO hPheo1 cells were used to create xenografts in SCID mice. *SDHB* KO cells were more likely to produce viable tumors than WT cells (Figure S3C). Tumors derived from *SDHB* KO cells exhibited increased calpain activity (Figure S3D) and exhibited higher levels of p35 and p25, while Cdk5 levels were comparable in both WT and KO tumors (Figure 2J). Tumor sections from both WT and *SDHB* KO xenografts immunostained for tyrosine hydroxylase (TH), and chromogranin A (CgA), considered as hallmarks of human neuroendocrine PC (Figure S3E) (Fliedner et al., 2010). Although not statistically significant, there was also a trend toward higher TH and ChrA levels in KO mice as well (Figure S3F). Together these data show that a shallow deletion of *SDHB*, similar to which occurs in humans, alters the metabolic profile, induces succinate buildup, and perturbs Ca^2+^/calpain/Cdk5 signaling in PC cells (Figure 2K and S3G).

### SDHB and Cdk5 correlation in human PC

Aberrant Cdk5 has been previously implicated in human neuroendocrine (NE) cancers (Pozo and Bibb, 2016). However, the link between Cdk5 and *SDHx* related NE tumors has heretofore not been investigated. Given that *SDHB* loss elicits elevated p35/25 levels in human PC cells, we assessed interactions between p35 mRNA and protein levels in human PC tissues. Interestingly, SDHB and Cdk5R1 (*i.e.,* encodes p35) were differentially expressed between PCs and adjacent normal adrenal medulla in TCGA and GEO datasets. A significant reduction in SDHB mRNA level (Figure 1D) corresponds to increased Cdk5R1 expression in PC compared to normal adrenals (Figure 3A). In addition, co-expression analysis of 161 PCPGs highlighted a significant negative correlation between SDHB and Cdk5R1 (R=-0.41, P=6.78 x 10^-8^, Figure 3B). No such significant correlation was observed between Cdk5R1 and other SDH subunit genes (such as SDHA, SDHC, SDHD). Also, no such correlation occurred between Cdk5R1 and other tumor suppressive genes of the TCA cycle (*e.g.*, *FH, IDH1, CS*) (Figure S4A-F). Furthermore, a PCPG GEO dataset of 84 patients showed a significant inverse correlation between expression of Cdk5 activating components (Cdk5, Cdk5R1 Cdk5R2) and SDHB (p<0.001) while no such relationships were observed with SDHA, SDHC and SDHD subunits (Figure S5A-C).

**Figure 3.**
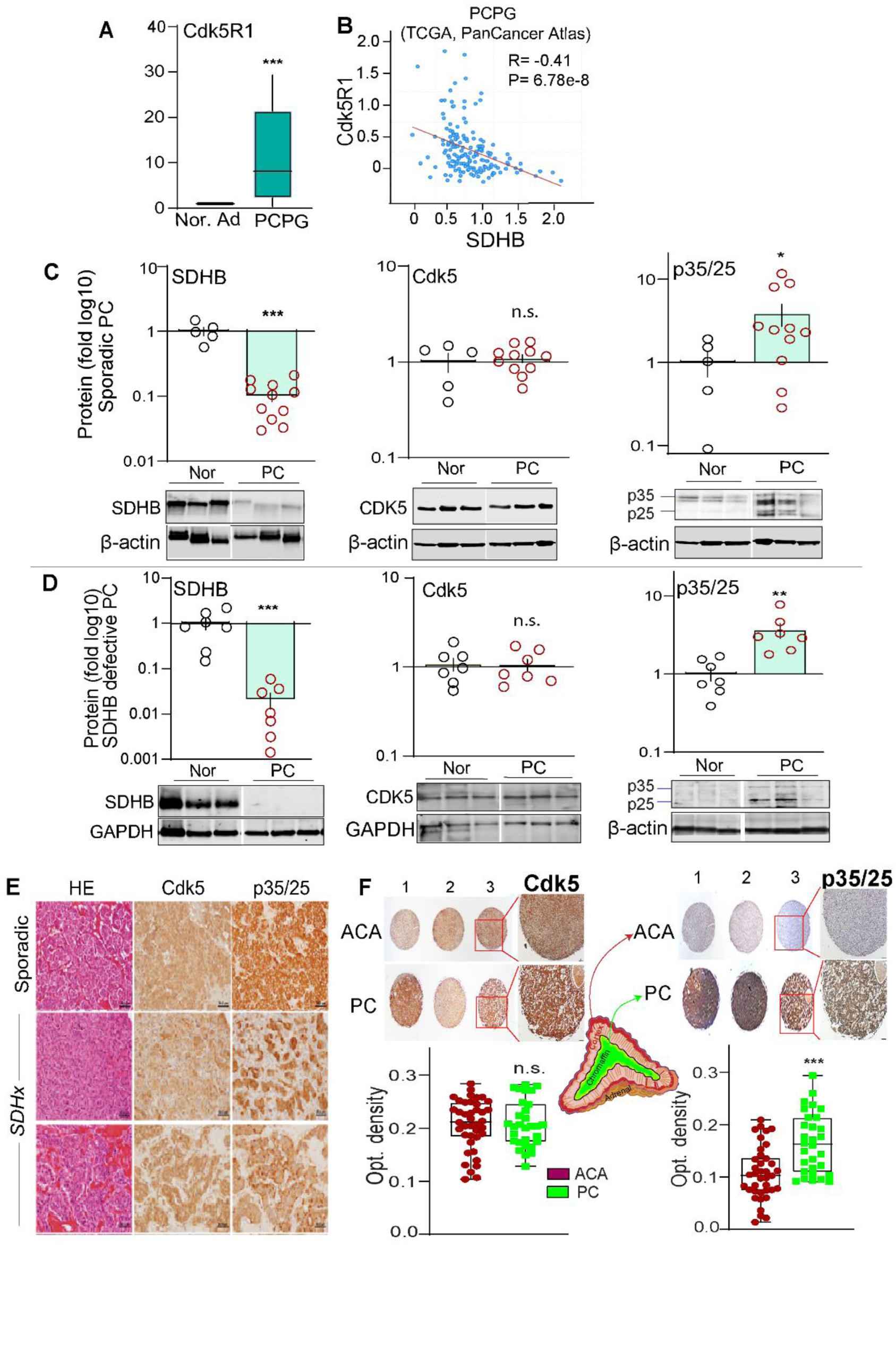
SDHB and Cdk5 coactivator expression inversely correlate in human PC. (A) Box plots of Cdk5R1 (*i.e.*, p35) expression in PC/PG tumors (n=179) vs. normal adrenal (n=3). (B) Correlation between SDHB and Cdk5R1 (p35), data quarried from cBioportal PCPG PanCancer atlas; (R is Spearman coefficient -0.41; p=6.78e). (C) Quantitation and representative blots showing protein levels of SDHB, Cdk5, and p35/25 in human sporadic PC tumors (n=11) compared with normal medulla (n=5). (D) Expression analysis of SDHB, Cdk5, p35/25 in human *SDHB* mutant tumors (n=7) vs. normal medulla (n=7). (E) HE and immunostains of Cdk5 and p35/p25 in human PC, scale bar = 50 µm. (F) Histological assessment of Cdk5, and p35/25 in tissue microarray sections of adrenocortical adenoma (ACA, n=40), and PC ( n=30); quantification presented as optical density; *p<0.05, **p < 0.01, ***p < 0.001 compared using *t*-test with Welch’s correction; n.s., non-significant.

Germline mutations in *SDHx* genes are responsible for 38-80% of metastatic and familial PCPGs, whereas approximately ∼10% of sporadic tumors also reportedly harbored *SDHx* mutations (Gottlieb and Tomlinson, 2005; Korpershoek et al., 2011). To determine if the inverse correlation between SDHB and p35 gene expression characterizes PC tumors at the post-translation level, immunoblots were performed with sporadic PC tumors vs. human normal adrenal medulla (Figure 3C). Consistent with the mRNA levels, SDHB protein levels decreased whereas p35 and p25 cumulative protein band intensities were increased (Figure 3C). This effect was mirrored in PC in which *SDHB* mutations were observed (Figure 3D). Notably, simultaneous increases of p35 (consistent with Cdk5R1 mRNA) and p25 in majority of PC tissues indicate the necessity to maintain sustainable protein levels of p35 to generate calpain cleaved p25.

Both sporadic and *SDHx* mutated PCs immunostained for Cdk5 and p35/p25 (Figure 3E). To determine cell type selectivity, differential expression of p35/25 was determined between PC tissues and a different form of adrenal tumor derived from cortical cells known as adrenocortical adenoma (ACA). Analysis of a human adrenal tissue microarray (30 cases of PC, and 40 cases of ACA) showed significantly higher levels of p35/25 in PC compared to ACA (1.5-fold, p = 0.0002) with no measurable change in Cdk5 (Figure 3F), implicating selective functional significance for aberrant Cdk5 in chromaffin cell-derived tumors. Taken together, these results illustrate that human sporadic and *SDHB* mutant PC manifest a negative correlation between SDHB and p35/p25, supporting the concept that aberrant Cdk5 can function as a strong driver of tumor cell proliferation in PC.

### Aberrant activation of Cdk5 in mouse chromaffin cells causes human-like PC

To determine if aberrant Cdk5 activation recapitulates human PC, a PiggyBac transgenic mouse carrying single copy PNMT transgene promoter was designed to drive cell-specific tetracycline transactivator (tTA) expression (Goldstein et al., 1972; Ross et al., 1990) (Figure 4A and S6A). This animal was then crossed with the Tet-Op-p25GFP line (Bujard, 1999) to generate bi-transgenic mice in which p25 overexpression (p25OE) could be induced in adrenal medulla chromaffin cells by removal of the tetracycline analog, doxycycline, from drinking water (Figure S6B) (Cruz et al., 2003). P25GFP expression status is denoted as p25-ON (Dox OFF) or p25- OFF (Dox-ON), and PNMT-tTA littermate controls are referred to as wild types (WT). Aberrant Cdk5 induced via activation of p25OE (p25-ON) caused bilateral 12-50 mm^3^ PCs to develop in 20-21 weeks with no effect on adrenal glands of mice lacking the Tet-Op-p25GFP transgene (WT) (Figure 4B and 4C).

**Figure 4.**
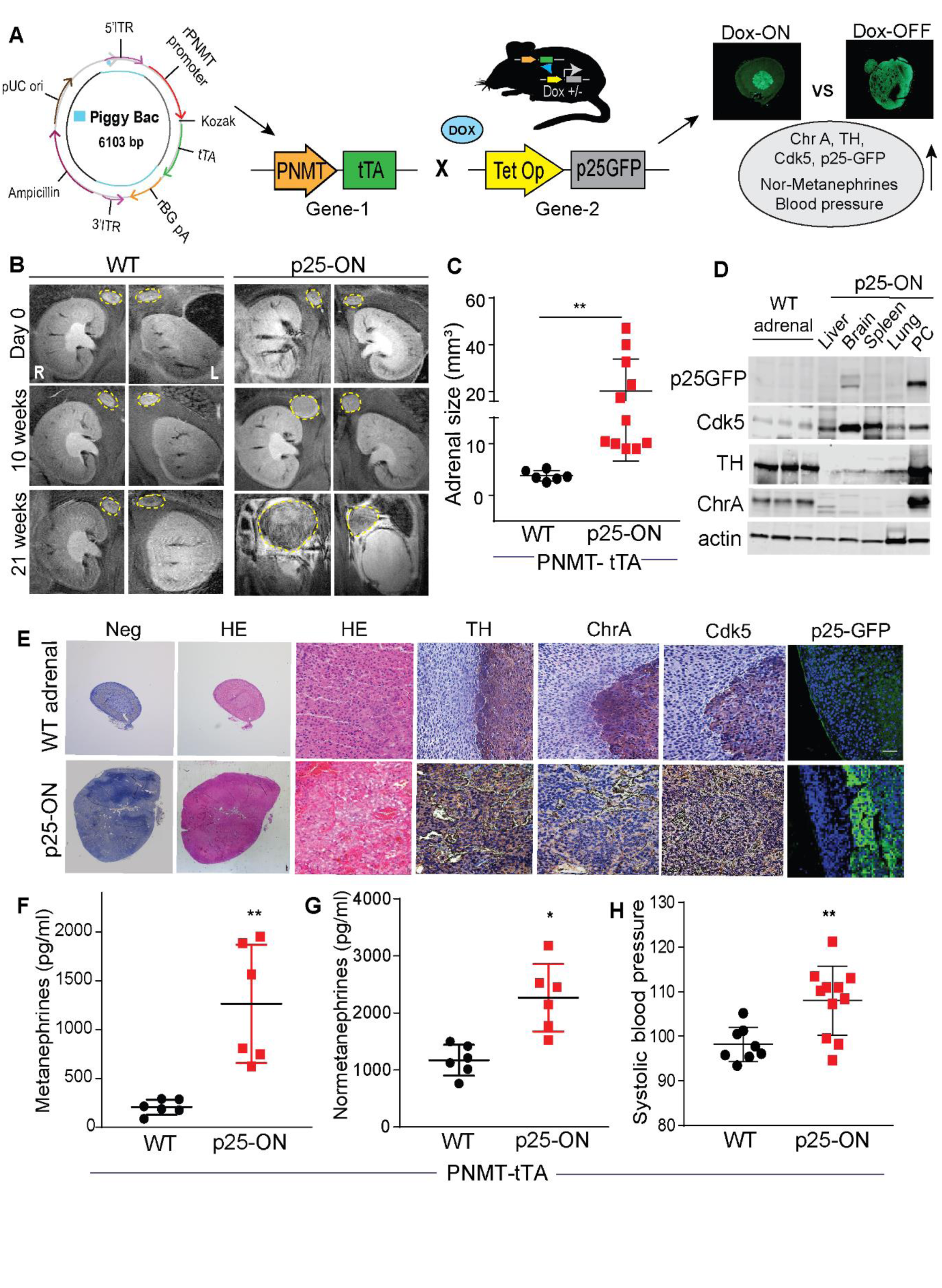
Aberrant Cdk5 develops PC in bi-transgenic mice. (A) Schematic of PiggyBac expression system designed to engineer single-copy PNMT–tTA (gene-1) crossed with Tet-Op- p25GFP (gene-2) to generate Dox ON/OFF bi-transgenic model. (B) T2-weighted representative MRI coronal images indicating temporal changes in adrenal gland size in WT (PNMT-tTA) vs. p25-ON (PNMT-tTA×p25GFP) mice. (C) MRI quantitation of adrenal size in p25-ON mice (n=11) compared to WT (n=6). (D) Immunoblots showing expression of p25-GFP, Cdk5, TH, and ChrA in tissue lysates derived from p25-ON or WT adrenal glands as indicated. (E) Histological assessment of NE markers in WT adrenals or PC tissue sections derived from p25-ON mice, scale bar 100 µM. (F-H) Measurements of plasma metanephrines (F), nor-metanephrines (G) and tail-cuff blood pressure (H); (n=6-11), *p < 0.05, **p < 0.01 compared by using *t*-test with Welch’s correction.

Immunoblots show higher levels of p25GFP, TH, and ChrA in p25-ON PC tissue compared to WT adrenal gland (Figure 4D). Moreover, p25-ON PC expressed the highest levels of these proteins compared to other organs including liver, brain, spleen, and lung, demonstrating tissue type specificity. Histological assessment of p25-ON tumor tissue showed neuroendocrine pseudo-rosettes surrounding blood vessels immunostained for TH/ChrA that typify human PC (Figure 4E). While both WT adrenal medulla and p25-ON PC stained for Cdk5, only p25ON tumors were p25-GFP positive (Figure 4E).

Human patients with PC have hypertensive crises due to tumor-derived overproduction of catecholamines (Eisenhofer et al., 1999). Similarly, p25-ON mice exhibited elevated levels of plasma metanephrine (Fig. 4F), normetanephrine (Figure 4G), and higher systolic blood pressure (Figure 4H) closely mirroring the human PC disease biochemical phenotype. These data demonstrate that aberrant Cdk5 activity, such as that resulting from *SDHB* mutations, can drive the formation and progression of PC and the clinical symptoms with which it is associated in a mouse model.

### Screening functional targets of aberrant Cdk5

Loss or gain of function mutations in kinases dysregulate signaling pathways associated with cancer (Hanahan and Weinberg, 2000). To understand how the aberrant Cdk5 invokes PC tumorigenesis, we analyzed our published library of phosphorylation sites derived from p25- ON/OFF NE tumors (Carter et al., 2020). Of 2000 proline-directed phosphorylation sites detected,

200 sites were upregulated, while 122 were downregulated, suggesting that p25OE both positively and negatively regulates protein phosphorylation in growing vs. arrested tumors (Figure 5A). To gain a wider perspective of phospho-dynamics we focused on phosphorylation sites suppressed in p25-ON tumors. Forty-four phosphoproteins were downregulated by >60% in growing tumors (Figure 5B). Functional analysis of these phosphoproteins revealed a network of enriched pathways involved in metabolic processes, cell cycle regulation, cell size, kinase activity, protein modification, and RNA processing (Figures 5C, S7, S8A-B).

**Figure 5.**
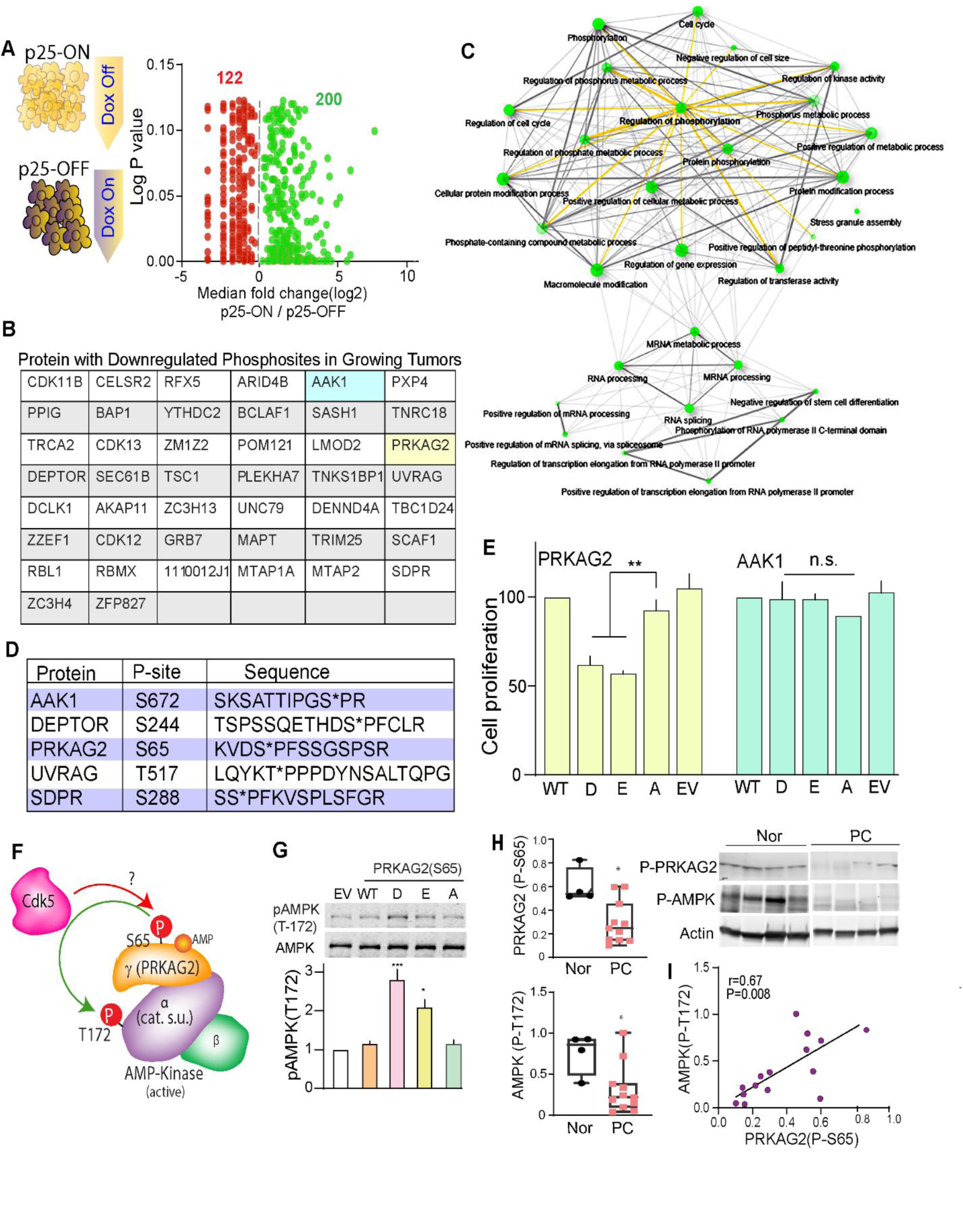
Characterization of functional Cdk5 targets in human PC. (A) Schematic showing NE tumors from p25-ON/OFF model analyzed for potential Cdk5 phosphorylation sites in PC. Function of phosphosite ratio (log2 fold change) between growing (p25-ON) and arrested (p25- OFF) tumors is plotted against p values. (B) Table showing Cdk5 phosphotargets downregulated in growing p25-ON tumors. (C) Graphical presentation of network tree highlighting enriched pathways associated with downregulated Cdk5 targets. (D) Table summarizing novel phosphosites selected to evaluate their effects on PC cellular proliferation. (E) Effects of phospho- and dephospho-mimetics (D/E/A) on cell growth was determined by dual fluorescence acridine orange/propidium iodide (AO/PI) viability staining, WT-wild-type; EV-empty vector. (F) Schematic of AMP-activated protein kinase heterotrimeric complex composed of α-, β- and γ-subunits. Novel phosphorylation site, Ser65 (S65) is located in PRKAG2 subunit of trimeric complex. (G) Immunoblot and quantification of phospho-Thr172 (P-T172) AMPK catalytic phosphorylation in response to PRKAG2 S65D/E/A phosphosite mutant expression, n=4. Values are means ± SEM, *p<0.05, **p<0.01, ***p<0.001, n.s. non-significant, one-way ANOVA multiple comparisons with Tukey’s method. (H) Quantitative immunoblotting of P-S65 PRKAG2 and -T172 AMPKα in human PC (n=11) vs. Normal adrenal medulla (n=4). Values are means ± SEM, *p<0.05, Student’s *t*-test. (I) Correlation plot between P-S65 PRKAG2 and -T172 AMPK in PC patients; r is Spearman’s correlation coefficient, **p<0.01.

To analyze phosphorylation sites of particular interest in PC, five targets were chosen based on the algorithm of having– 1. Novel phosphorylation sites; 2. Conserved phosphosite in humans; 3. Implicated in cancer metabolism, protein trafficking, cell cycle, autophagy, or protein translation.

These phospho-targets included; phospho-Ser672 AAK-1, phospho-Ser244 DEPTOR, phospho- Ser65 PRKAG2, phospho-Thr517 UVRAG, and phospho-Ser288 SDPR (Figure 5D). To assess functional relevance to PC tumorigenesis, each phosphorylation site was mutated to encode phospho-mimetic (D/E) or phospho-null (A) amino acids and transiently overexpressed in hPheo1 cells. Of the five phospho-targets, S65D/E forms of PRKAG2 significantly suppressed cell proliferation compared to that of WT or the S65A mutant (Figure 5E).

PRKAG2 is the non-catalytic regulatory γ2 subunit of AMP-activated protein kinase (AMPK), a heterotrimeric metabolic regulator composed of catalytic α-subunit and two regulatory (β and γ) subunits (Figure 5F). AMPK regulates cellular energy homeostasis, glucose sensing, and immune response processes that impact cell growth and proliferation (Zadra et al., 2015). PRKAG2 has been chiefly studied in cardiac and skeletal muscles (Pinter et al., 2013; Zhan et al., 2018). However, the isoform-specific function of PRKAG2 is largely unexplored in cancer. Nucleotide binding to PRKAG2 confers phosphorylation/dephosphorylation upon residue Thr172 in the activation loop of the AMPK catalytic α-subunit, as a prerequisite to kinase activation (Gowans et al., 2013; Oakhill et al., 2011; Shaw et al., 2004). Interestingly, D/E mutations of S65 PRKAG2 increased phospho-Thr172 AMPKα (Figure 5G and S8C). This suggests that phosphorylation at Ser65 PRKAG2 controls AMPK activity. These data also implicate the regulation of AMPK via phospho-Ser65 PRKAG2 as an important downstream effector of aberrant Cdk5 in NE cells.

### Aberrant Cdk5/GSK3 deregulates AMPK pathway

AMPK activity is regulated by either allosteric or non-canonical mechanisms (Hardie, 2014; Hawley et al., 2005). To better understand how Ser65 PRKAG2 phosphorylation could regulate AMPK activity, a phosphorylation state-specific antibody was generated to this site (Figure S9A- C). Quantitative immunoblot analysis of human patient tissue showed that PC tumors had significant decreases in both phospho-Ser65 PRKAG2 and -Thr172 AMPKα compared to normal control adrenals, with a linear correlation between the two sites (Figure 5H-I). In contrast, no alterations in PRKAG2/AMPK gene expression were observed between normal and tumor tissues (TCGA-GEPIA dataset; n=182, Figure S9D). Thus decreased Ser65 phosphorylation and AMPK inactivation characterizes both human and mouse NE tumors.

A decrease in phospho-Ser65 PRKAG2 and AMPK activity in response to aberrant activation of Cdk5 suggests an intermediary signaling step is involved. In fact, both Cdk5 and GSK3 are predicted to phosphorylate Ser65 PRKAG2 (Fig. S9E). Since Cdk5 activation indirectly results in GSK3 inactivation (Morfini et al., 2004), we asked if Ser65 PRKAG2 phosphorylation by GSK3 is downstream of aberrant Cdk5/p25. Indeed, GSK3 efficiently phosphorylated purified recombinant PRKAG2 (Figure 6A). The site at which GSK3 phosphorylates PRKAG2 was confirmed as Ser65 by immunoblotting, while Cdk5 did not phosphorylate this site *in vitro* (Figure 6B-C). Also, active GSK3 immunoprecipitated from hPheo1 cell lysates efficiently phosphorylated WT PRKAG2 *in vitro* compared to the S65A recombinant protein (Figure S9F-G). Aberrant Cdk5 activation leads to inhibitory phosphorylation of Ser9 GSK3β (Plattner et al., 2006; Wen et al., 2008). In agreement, *SDHB* KO PC cells exhibited higher phospho-Ser9 GSK3β levels compared to parent cells (Figure S9H). Also, expression of kinase-dead (KD) vs. WT Cdk5 reduced phospho-Ser9 GSK3 by 50% and caused a concomitant increase in phospho-Ser65 PRKAG2 and Thr172 AMPKα (Figure 6D).

**Figure 6.**
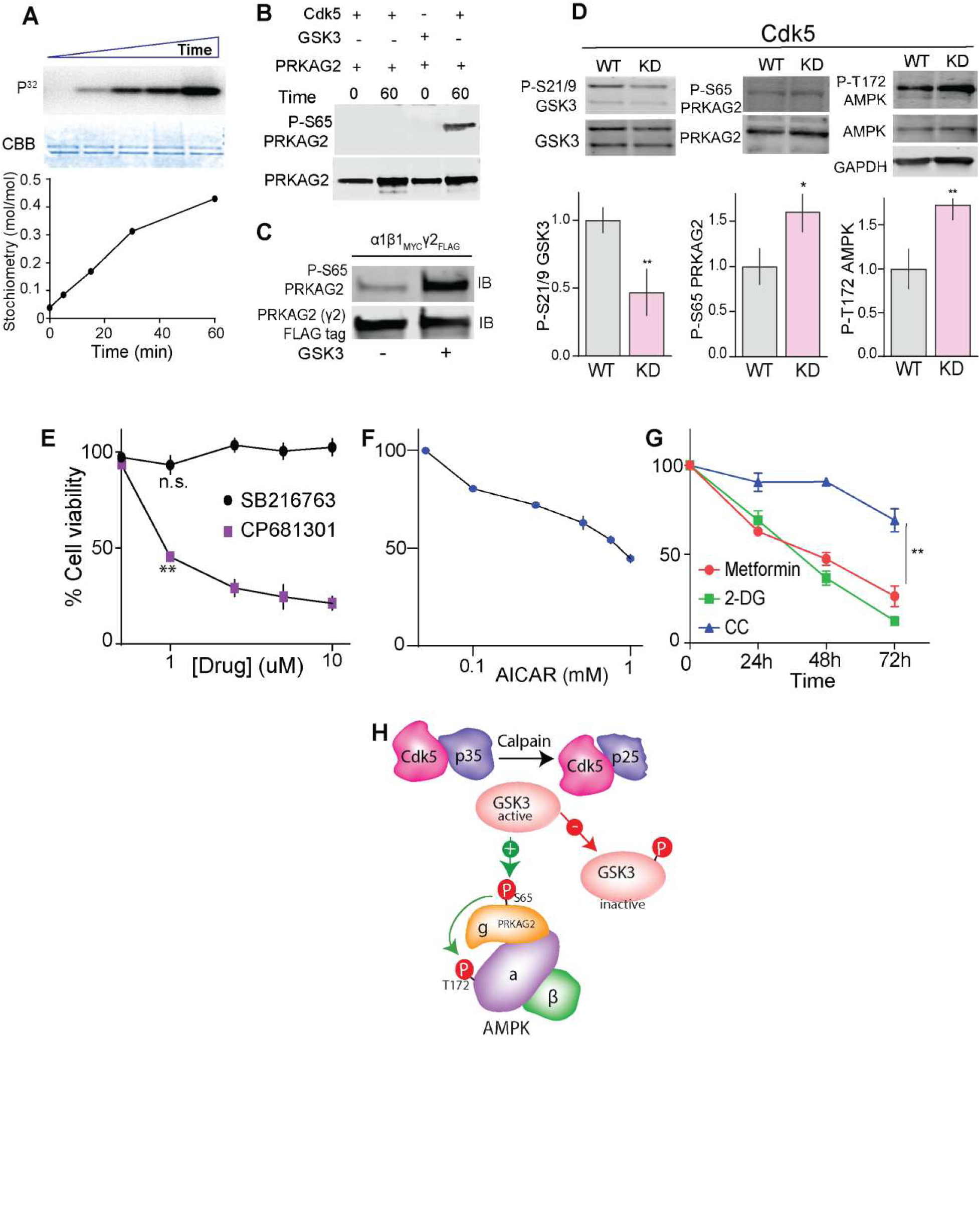
Cdk5/GSK3 signaling regulates P-PRKAG2/P-AMPK. (A) *In vitro* phosphorylation of recombinant PRKAG2 by GSK3, with time-dependent ^32^P incorporation and Coomassie stained protein (CBB) shown with stoichiometry. (B) Immunoblots showing GSK3 but not Cdk5 phosphorylates Ser65 PRKAG2 *in vitro*. (C) Immunoblots of recombinant AMPK holoenzyme trimeric complex (α1β1γ2) phosphorylated by GSK3β. (D) Effects of ectopic expression of wildtype (WT) vs. kinase dead (KD) Cdk5 vector on the levels of indicated phosphorylation sites in PC cells. (E) Dose-dependent effects of GSK3 (SB216763, 24 h) vs. CDK5 inhibition (CP681301, 24 h) on *SDHB* KO hPheo1 cell viability. (F) Plot showing dose dependent effect of AMPK activator AICAR on cell viability treated for 24 h. (G) Time-dependent effects of AMPK activators, Metformin (20 mM), 2-Deoxy-D-glucose (2-DG, 20 mM), and AMPK inhibitor, Compound C (10 μM) on KO hPheo1 cell viability. n=3, values are mean ± SEM, *p<0.05, \*\**p* < 0.01 compared by one-way ANOVA. (H) Schematic of the potential signaling mechanism where aberrant CDK5/p25/GSK3β crosstalk regulates phospho-dynamics between S-65 PRKAG2 and T-172 AMPKα.

GSK3 has been broadly implicated in cancer and inhibitors or activators of GSK3 may induce cellular proliferation or suppression dependent on cell type (Li et al., 2014; Pap and Cooper, 1998; Tang et al., 2012). Here, we found that hPheo1 cells were unaffected by treatment with a GSK3 inhibitor (SB216763), while a Cdk5 inhibitor (CP681301) dose-dependently abrogated cell viability (Figure 6E). Moreover, metabolic modulators which activate AMPK, including AICAR, metformin (Vial et al., 2019) and 2-DG (Wang et al., 2011) induced growth inhibitory effects whereas, an AMPK inhibitor, Compound C (dorsomorphin) (Zhou et al., 2001) had a minimal effect on cell growth (Figure 6F-G). These results suggest a novel signaling cascade where aberrant Cdk5 activation causes inhibitory GSK3 phosphorylation, thereby disrupting phospho-PRKAG2- dependent AMPK activation (Figure 6H).

### Characterization of novel Cdk5 inhibitor

With the emergence of Cdk5 as a promising target in several cancers (Pozo and Bibb, 2016), there is renewed demand for effective drugs which target this kinase. Currently available Cdk5 inhibitors such as Roscovitine (CYC202, Seliciclib) act as purine analogs that interfere with ATP binding (Bettayeb et al., 2010). However, their lack of specificity, short half-life, rapid degradation, and weak potency limit their potential for clinical use (McClue and Stuart, 2008). Therefore, we screened a small library of selective inhibitors of Cdk5, which included 25-16, MRT3-007, MRT3- 124, and CR8 using Roscovitine as a positive control. These compounds share the same general chemical structures (shown for CR8, MRT3-007, Roscovitine, Figure 7A) and relative kinase selectivity (Figure 7B). Each of these compounds exhibited dose-dependent effects on hPheo1 cell viability (Figure 7C). MRT3-007 showed highest potency with an IC_50_ value approximately 1000-fold lower than that of Roscovitine (25 ± 10 nM for MRT3-007 vs. 26 ± 10 µM for Roscovitine).

**Figure 7.**
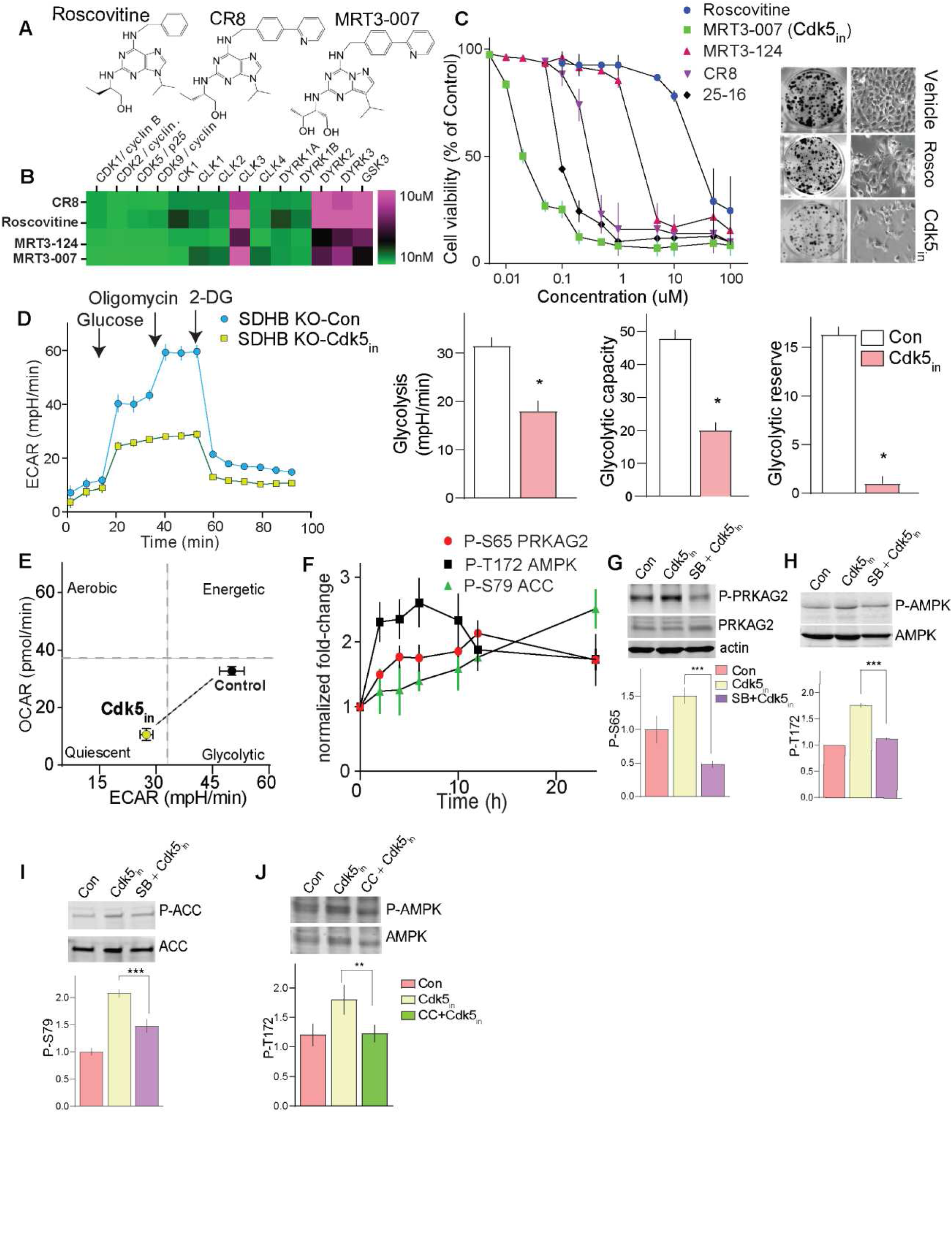
Targeting Cdk5 to regulate the P-PRKAG2/P-AMPK cascade. (A) Chemical structures of Cdk5 inhibitors, Roscovitine, CR8 and MRT3-007. (B) IC50 heatmap showing *in vitro* kinase profile of CR8, Roscovitine, MRT3-124, and MRT3-007. (C) Dose-response effects of Cdk5 inhibitors on cell viability (left) and colony formation (right) of hPheo1. n=4, data are means ± S.E.M. (D) Glycolytic profile of *SDHB* KO cells treated with or without MRT3-007 ([Cdk5_in_], 25 nM for 12 h, (left). Plots comparing basal glycolysis rate, glycolytic capacity, and glycolytic reserve between control versus MRT3-007 (Cdk5_in_) (right). (E) Bioenergetic phenotype of KO cells in response to MRT3-007. Data are means ± SD, *p < 0.05, Student *t*-test, n=2 (10 replicates per group). (F) Immunoblot quantification showing time dependent effect of MRT3-007 (Cdk5_in_) on phosphorylation states of P-S65 PRKAG2, P-T172 AMPK and P-S79 ACC respectively. n=3, values presented as fold change normalized with time=0. Data are mean ± S.E,M. (G-I) Immunoblot and densitometric analysis presents impact of GSK3 inhibitor, SB216763 (5 µM, pretreatment 10 h) on Cdk5_in_-induced phosphorylations as indicated. (J) hPheo1 pre-incubated with or without compound C (CC, 10 µM), representative blot and quantitation comparing effects of Cdk5_in_ alone or in combination with CC as indicated.

Effective cancer treatments using kinase inhibitors depend on precise genetic constitution of individual patients so that differences in molecular signatures between tumor and normal cells can be defined (Broekman et al., 2011; McDermott et al., 2007). Furthermore, it is possible for inhibitors to be effective by targeting the kinase driving neoplasia while exhibiting broader activity *in vitro*. As a typical example, MRT3-007 shares overlapping selectivity for Cdk1, 2 and 9 *in vitro*. Therefore, we queried whether its growth inhibitory effects in PC were Cdk5 dependent. Notably, PCPG patients showed significantly higher gene expression of Cdk5 over Cdk1/2 (Figure S10A), while Cdk9 expression was absent in the experimental models used in this study. Consistent with this, MRT3-007 was more effective in suppressing *in vitro* growth of PC compared to selective inhibitor of Cdk2 (CVT-313) (Figure S10B). Also, MRT3-007 elicited limited efficacy on cancer cell lines which generate lower p25 levels (such as those derived from breast cancer (MDAMB231 cells), liver carcinoma (HepG2 cells), and small cell lung cancer (H1184 cells). In contrast, MRT3- 007 showed higher potency for cervical cancer cells (HeLa cells), which express higher p25 levels, comparable to that observed in PC-derived cells (Figure S10C). These analyses suggest that this class of kinase inhibitor could be effective in treating tumors that are dependent upon hyperactive Cdk5 compared to other off target kinases.

Additional support for *ex vivo* selectivity of MRT3-007 for Cdk5 over Cdk1/2 was derived by immunoprecipitating Cdk5 or Cdk1/2 from the lysates of hPheo1 cells treated with MRT3-007 using histone H1 as a reporter substrate. Cdk5 but not Cdk1/2-dependent phosphorylation of histone H1 was significantly attenuated, suggesting MRT3-007 is more selective for Cdk5 within the contest of the intracellular milieu than *in vitro* (Figure S10D). MRT3-007 also selectively inhibited the proliferation of cells that overexpress p25 (p25OE, Dox Off) but had no effect on cells lacking p25 expression (Dox On, Figure S10E) (Pozo et al., 2013), implying aberrant Cdk5- dependent sensitivity. Together these results suggest that MRT3-007 can be exploited as a potent Cdk5 inhibitor (Cdk5_in_) in cells, which depend upon aberrant Cdk5 for their growth.

### Cdk5-GSK3-AMPK cascade controls bioenergetics and induces senescence

Cdk5 plays a critical role in tumor-associated cell cycle progression, DNA damage response, and mitochondrial dysfunction (Mao and Hinds, 2010; Sun et al., 2008; Zhang et al., 2015). Here, we showed that loss of *SDHB* in hPheo1 cells caused a metabolic shift towards glycolysis (see Figure 1F) and triggered aberrant Cdk5-AMPK signaling to drive cell proliferation. These findings prompted us to explore the impact of novel Cdk5 inhibitor on AMPK signaling cascade, downstream metabolism, and cell cycle progression. Treatment of *SDHB* KO cells with Cdk5 inhibitor– MRT3-007 (Cdk5_in_), decreased basal glycolysis, glycolytic capacity and glycolytic reserve in response to sequential addition of glucose, oligomycin and 2-DG (Figure 7D). In addition, Cdk5_in_ shifted the cell energy profile (ratio of OCR:ECAR) from a glycolytic to quiescence/low energy state (Figure 7E). Concomitantly, Cdk5 inhibition caused time-dependent increases in phopsho-S65 PRKAG2 and -T172 AMPKα (Figure 7F). These effects corresponded to increased phosphorylation of S79 acetyl-CoA carboxylase (ACC), a defined AMPK activity reporter, confirming that Cdk5_in_ activates the AMPK pathway.

To better understand the novel Cdk5-GSK3-AMPK signaling pathway, the GSK3 selective inhibitor, SB216763, was used to determine the role of GSK3 in Cdk5_in_-mediated AMPK activation. Pretreatment with SB216763 suppressed activation of S65-PRKAG2, T172-AMPK, and S79-ACC phosphorylation by Cdk5_in_ (Figure 7G-I). Furthermore, direct inhibition of AMPK via compound C (CC) reversed the effects of Cdk5_in_ on phospho-T172 AMPK (Figure 7J). These data further substantiate the Cdk5-GSK3-AMPK cascade as a critical signaling pathway downstream of SDHB deficiency that mediates neoplastic cell proliferation.

On sensing bioenergetics stress, AMPK functions as a checkpoint and regulates cell cycle machinery via phosphorylation of the tumor suppressor protein, p53. AMPK-dependent phosphorylation of Ser15 p53 stabilizes the protein and is essential to cell cycle arrest (Jones et al., 2005) (Garcia and Shaw, 2017). In agreement, Cdk5_in_ caused a time-dependent increase in phospho-Ser15 p53 (Figure 8A), implicating p53 as a critical downstream effector of AMPK in hPheo1 cells. P53 functions as a transcription factor controlling the expression of cell cycle or growth-regulating genes, (Chen, 2016; Mijit et al., 2020) while AMPK-p53 activation can induce cellular senescence in response to bioenergetics stress or chemotherapy (Jones et al., 2005; Lee et al., 2015; Xue et al., 2007). Consistent with this, in addition to increased Ser15 p53 phosphorylation, Cdk5_in_ induced time-dependent increases in the expression of senescence markers, p16^INK4a^ and p27^Kip^, which associate with the primary G1-S checkpoint (Figure 8B-C). Cdk5_in_ also caused a sharp rise in phospho-Ser139 histone H2Ax, a marker of DNA damage such as that which occurs in response to chemotherapy (Figure 8D) (Zhao et al., 2019). These effects in response to Cdk5_in_ were negated in the presence of the p53-specific inhibitor, Pifthrin (Pftα) (Figure S11A). Furthermore, Cdk5_in_ induced senescence-like cell morphology, marked by enlarged flattened cells (Figure 8E), disorganized cytoskeleton, and elevated β-galactosidase, concomitant with overexpression of p16^INK4a^, p27, and amplification of phospho-S139 H2Ax foci (Figure 8F). Of note, inactivation of AMPK via compound C reversed the cell spreading induced by Cdk5_in_, indicating that AMPK activation mediated the Cdk5_in_-induced morphological changes (Figure S11B).

**Figure 8.**
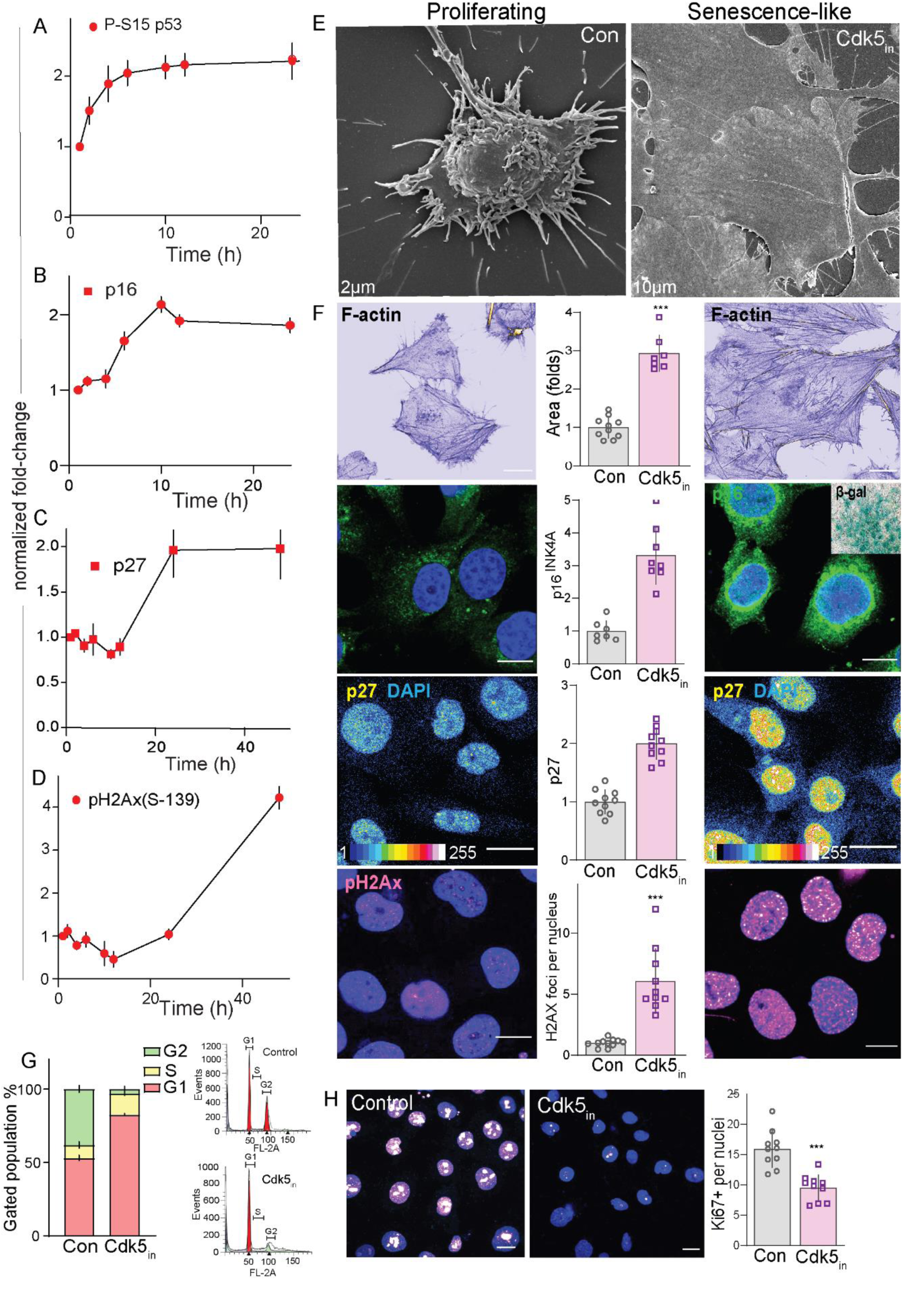
Cdk5_in_ induces senescence–like phenotypic characteristics. Immunoblots quantification showing time dependent effect of Cdk5_in_ (*i.e* MRT3-007) on phosphorylation states or levels of (A) P-S15 p53, (B) p16^INK4a^,(C) p27^Kip^, (D) P-S139 H2AX. Values presented as fold change normalized with time=0. n=3, data are means ± S.E,M. (E) Scanning electron microscopy (SEM) images of hPheo1 cell morphology in control vs. Cdk5_in_ (Indolinone A). (F) Imaging of proliferating cells and those treated with Cdk5_in_ (MRT3-007, 20 nM, 48 h) for common senescent markers. Representative confocal photomicrographs and quantitation shows phalloidin stain of F-actin, p16^INK4a^ (inset is senescence-associated-β-galactosidase; β-gal), p27^Kip^ and P-H2Ax (S139) respectively; scale=136 µM. (G) Representative DNA histogram (left) showing cell cycle profile of KO cells treated with vehicle or Cdk5_in_ (20 nM); histograms analysis by MODfit LT 3.0 (right) presents cell cycle distribution in G1 (red), early-S (yellow), and late-S/G2/M (green); n=2, Data are means ± SD. (H) Confocal images of Ki67 staining and bar graph show number of Ki67- positive cells.

The observed cell phenotypic effects of Cdk5 inhibition also corresponded to a significant shift toward cell cycle arrest in G1 phase with a drastic expulsion of cells from G2 phase (Figure 8G). At the same time, a decrease in proliferation marker Ki67 was evident (Figure 8H). The G1/S cell cycle arrest was also observed in cells that overexpressed S65D PRKAG2 compared to those expressing either WT or S65A forms of the AMPK regulatory subunit (Figure S11C). Together, these data show that Cdk5 inhibition not only alters the bioenergetics of cancer cells, but also provokes senescence-like characteristics following activation of PRKAG2/AMPK/p53 signaling cascade.

### Cdk5 inhibition as a preclinical treatment for SDHB-mediated disease

The above results implicate CDK5_in_ (*i.e.*, MRT3-007) as a promising targeted therapy for PC and other SDHB-related disorders. To evaluate its anti-tumor potential *in vivo,* the maximal tolerated dose (MTD) of CDK5_in_ was first determined in mice. MRT3-007 was well tolerated up to dose of 1 mg/kg (Figure S12A). Subsequently, mice carrying *SDHB* KO hPheo1 xenografts were treated with 0.5 mg/kg MRT3-007, which induced a significant reduction in tumor volume and mitotic index (measured by Ki67 expression, Figure 9A-B) with minimal adverse effect on body weight (Figure S12B). Additionally, *in vivo* therapeutic efficacy of CDK5_in_ was confirmed using an established metastatic allograft model of PC where luciferase-expressing mouse PC cells (MTT) were injected intravenously and metastases were imaged *in vivo* (Martiniova et al., 2009). CDK5_in_ dramatically reduced tumor signal compared to both vehicle and its parent compound, roscovitine (Figure 9C). Consistent with our *ex vivo* findings, Cdk5 inhibition reduced phospho-Ser21/9 and increased phospho-Ser65 PRKAG2, -Thr172 AMPK, and -Ser79 ACC in the lysates of PC xenografts (Figure 9D-G). Additionally, CDK5_in_ treatment increased phospho-Ser15 p53, p16, p27 levels, consistent with that of senescence-like markers observed *in vitro* (Figure 9H). These findings indicate that CDK5_in_ possesses therapeutic potential in the treatment of PC and other *SDHB*- related disorders.

**Figure 9.**
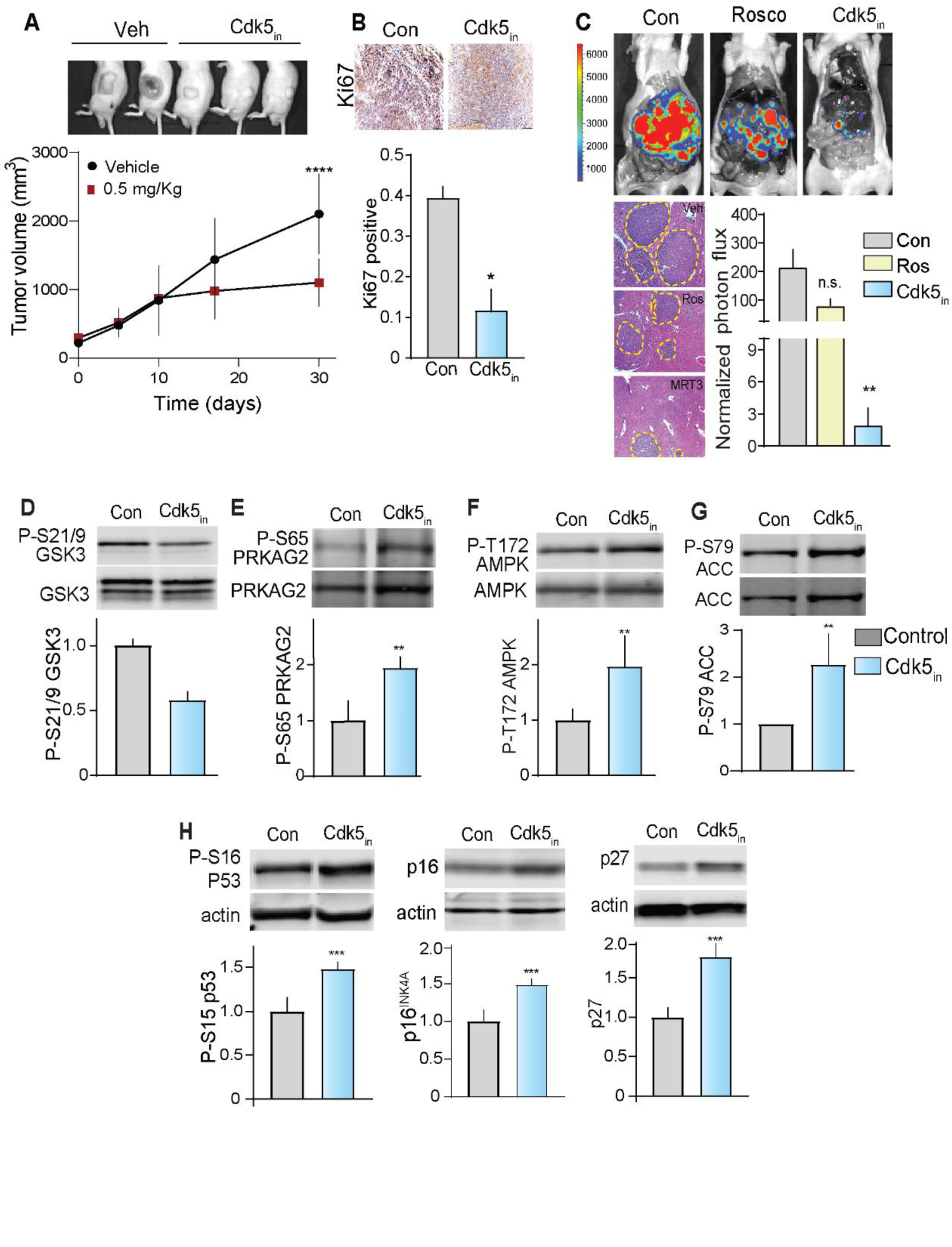
Cdk5 inhibition attenuates *in vivo* tumor growth. (A-B) Tumors in mice implanted with *SDHB* KO xenografts and treated with vehicle (C, n=8) or Cdk5_in_ (T, n=10) (0.5 mg/Kg) were analyzed for volume (A) and cell proliferation (Ki67, B). (C) Efficacy of Cdk5 inhibition tested in a metastatic allograft PC model, examined via *in vivo* bioluminescent imaging. Representative images of vivisected liver (top) and tumor spread indicated by histological analysis and photon flux quantitation (bottom). (D-H) Quantitative immunoblots of phosphorylation sites and proteins indicated in xenograft tumor lysates. n = 5-8, Data are means ± SEM, *p < 0.05, **p < 0.01, ***p < 0.001 compared by Student’s *t*-test.

Finally, to validate that PC tumor progression is dependent upon aberrant Cdk5 activity, we assessed the effects of halting p25OE in the bitransgenic mouse PC model. Under Dox-Off (p25- ON) conditions, tumor size progressed by 2.5-fold over 18 weeks. However, replacement of dietary Dox (Dox-On, p25-OFF) significantly limited tumor volume with a corresponding decrease in p25-GFP expression (Figure 10A-C). Tumor arrest due to halt in p25OE (p25-OFF) resulted in decreased phospho-Ser21/9 GSK3 and increased phospho-Ser65 PRKAG2, -Thr172 AMPK, - Ser79 ACC (Figure 10D-G). Once again, these tumors achieve a senescent-like state as indicated by increased phospho-Ser15 p53 and p16 levels (Figure 10H-I).

**Figure 10.**
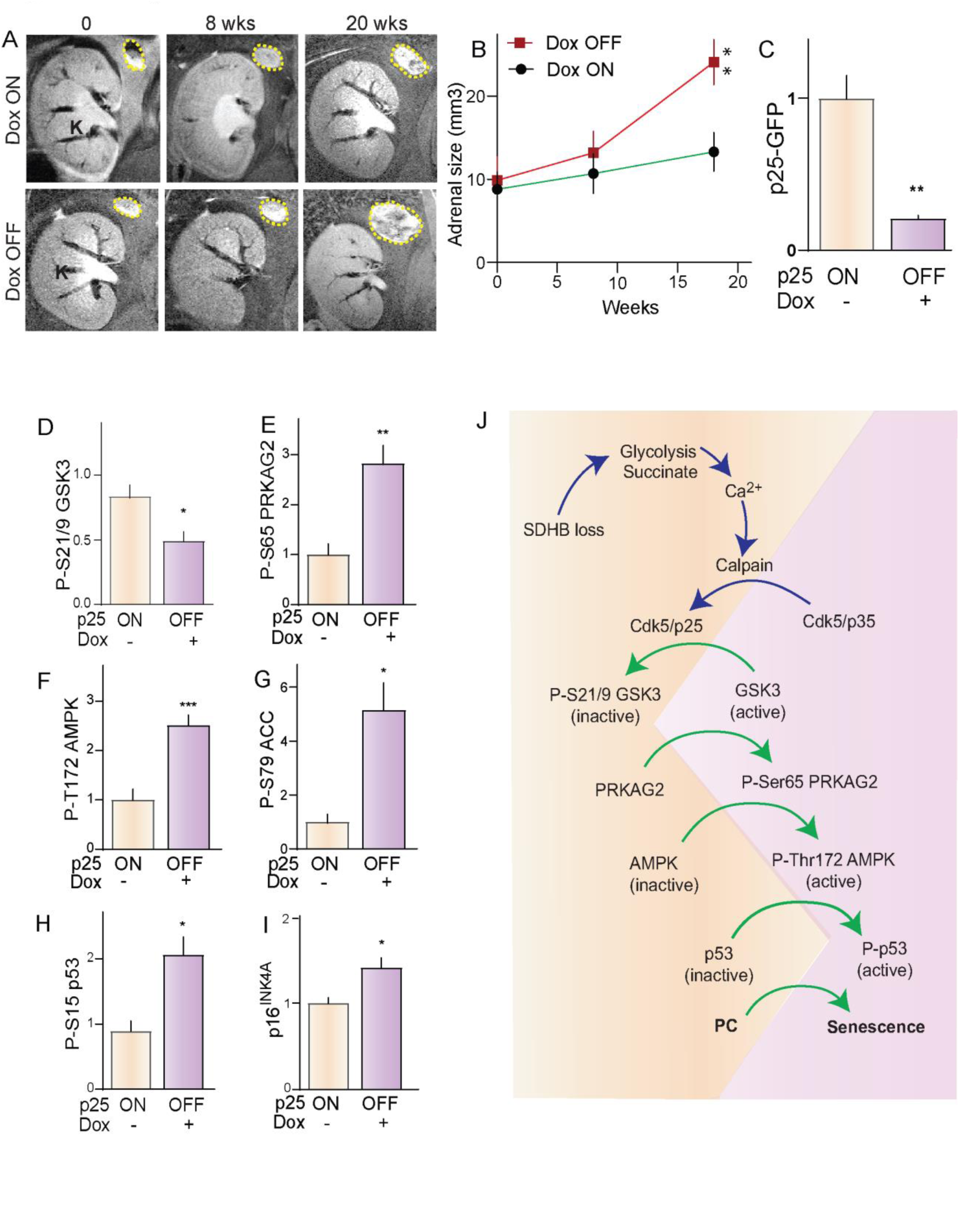
Halt of p25GFP expression arrests PC growth in bitransgenic mice. (A-B) Representative coronal MRI images and quantitation showing changes in adrenal gland size of mice over-time under Dox ON and OFF conditions. (C) Quantitation of p25GFP expression determined by immunoblotting. (D-I) Immunoblot quantitation of indicated phosphorylation site or protein from p25ON/OFF tumors derived from bitransgenic mice. Data are means ± SEM; \**p* < 0.05, \*\**p*<0.01, \*\*\**p*< 0.001 Student’s *t*-test, n=4-7. (J) Schematic depicting the signaling cascade delineated in these studies downstream of SDHB loss.

Together, these data support a signaling cascade (Figure 10J) where SDHB loss causes accumulation of succinate which in the context of metabolic impairment leads to loss of Ca^2+^ control and calpain activation. As a result, aberrant Cdk5/p25 accumulates and causes GSK3 inactivation. Consequently, AMPK is inactivated through the reduction in Ser65 PRKAG2 phosphorylation. Attenuated AMPK activity leads to reduced phosphorylation of Ser15 p53 promoting cell proliferation. This signaling cascade appears to be an important additional feature of the Warburg effect.

## DISCUSSION

Significant defects in any component of Krebs/TCA cycle is problematic because metabolite intermediates such as succinate act as key signals orchestrating metabolic reprograming inherently tied to the cell proliferative state (Zhao et al., 2017). TCA-linked mitochondrial malfunction and elevated glucose utilization is increasingly viewed as a root cause of several human diseases including cancer, diabetes, and neurodegenerative disorders. (Blank et al., 2010; Hsu and Sabatini, 2008). Here, we delineated a novel signaling cascade, which highlights critical phosphorylation hotspots on metabolic checkpoints disrupted by loss of the TCA component, *SDHB*. Succinate buildup stemming from loss of function of SDHx enzyme complex contributes to a plethora of pathophysiological processes. Previously, several downstream effects of succinate accumulation have been linked to altered metabolism, pseudohypoxic HIF1/2α stabilization, oxidative stress, SUCNR1 activation and tumor-associated inflammation (Dahia et al., 2005; Matlac et al., 2021; Pollard et al., 2006). Though these mechanisms have been demonstrated to serve as components in discrete cellular- or disease-specific contexts, their suggested interactions as part of the multistep process of carcinogenesis have not been fully delineated. Hence, the complete picture of how TCA perturbations can lead to cancer has not yet emerged.

Our study shows that activation of aberrant Cdk5 signaling in response to succinate buildup plays central role in regulating metabolic/cell cycle checkpoints. Previously, features of neurodegeneration or ischemic injury has been modeled by inhibiting SDH activity using 3-NPA (3-nitropropionic acid), and malonate, which in turn activates calpain/Cdk5 signaling (Barros- Minones et al., 2013; Pang et al., 2003; Ranganayaki et al., 2021). These findings independently support the importance of Cdk5 in *SDHx* deficiency diseases. Here we deciphered several distinct aspects of tumorigenic signaling including metabolic shift coupled with mismanagement of [Ca^2+^]_i_/calpain/aberrant Cdk5 in response to *SDHB* loss in PC tumors.

Excessive succinate freely shuttles across cell membrane via INDY transporters or between the mitochondria and cytosol via the dicarboxylic acid translocator/voltage-dependent anion channels and released into the extracellular milieu (de Castro Fonseca et al., 2016). The extracellular succinate trigger SUCNR1 receptor where invokes multiple signaling outcomes, dependent on the cell type, while excess intracellular succinate accumulation causes inhibition of 2-oxoglutarate-dependent dioxygenases (*e.g.*, prolyl hydroxylases), and histone and DNA demethylases. Here, we showed for the first time that aberrant Cdk5 acts as a downstream effector of SDHB-succinate-Ca^2+^ signaling. Succinate accumulation could have caused [Ca^2+^]_i._ dysregulation in hPheo1 cells either through activation of SUCNR1 receptors or mitochondrial ROS (Andrienko et al., 2017). However, we also observed elevation of SUCNR1 in hPheo1 cells which was consistent with previous studies where silencing *SDHB* induced SUCNR1 expression in HepG2, MTT cells (Cervera et al., 2008; Richter et al., 2018). Importantly, we showed that SUCNR1 inhibition prevented intracellular Ca^2+^ mobilization in *SDHB* KO cells, supporting the succinate-SUCNR1 pathway as a primary source of [Ca^2+^]_i._ dysregulation in PC cells.

While Cdk5 is recognized for functions in neurobiology, it has only recently emerged as an important factor in cancer (Antoniou et al., 2011; Carter et al., 2020; Dorand et al., 2016; Herzog et al., 2016; Pozo and Bibb, 2016; Pozo et al., 2013). The present study defines a new role for Cdk5 in cellular metabolism and PC etiology in particularly. We discovered that aberrant Cdk5 activation attenuated AMPK activity as cell metabolism shifts towards glycolysis in PC cells following *SDHB* loss. Furthermore, Cdk5-dependent GSK3 inhibition served as an intermediate step in AMPK regulation. Cdk5-GSK3β interactions have been demonstrated in neurodegenerative disorders although precisely how Cdk5 induces GSK3 inhibition remains to be delineated. This may involve either activation of ErbB/Akt (Wen et al., 2008) or inhibition of phosphatases such PP1/PP2A (Morfini et al., 2004; Plattner et al., 2006) that increase inhibitory Ser9 GSK3 phosphorylation in response to Cdk5 activation. The data presented here strongly support Cdk5 as the arbitrator regulating the phospho-dynamics of a GSK3/PRKAG2/AMPK cascade which determines the proliferative state of PC cells.

While we showed that GSK3 can activate AMPK via PRKAG2 phosphorylation in PC cells, it has also been suggested that GSK3 interacts with the AMPKβ regulatory subunit and can inhibit AMPK by phosphorylating Thr479 of the catalytic subunit (Suzuki et al., 2013). This signaling mechanism is somewhat confounded as other studies demonstrated that activation of GSK3/AMPK is required for regulation of protein synthesis and cellular proliferation (Jorgensen et al., 2004; Park et al., 2014). Also, AMPK activators such as phenorphin and AICAR has been shown to activate AMPK in parallel with activation of GSK3 by alleviating inhibitory S9 phosphorylation in neuronal models (King et al., 2006). Thus, the relationship between GSK3 and AMPK may be context-dependent.

Canonical regulation of AMPK involves binding of AMP or ADP to γ-regulatory subunits (PRKAG-1,2,3), resulting in phosphorylation of Thr172, located in the kinase catalytic subunit activation loop, by LKB1. Non-canonical regulation of AMPK is chiefly characterized by Thr172 phosphorylation by CAMKK2. We identified a functional role for Ser65 PRKAG2 phosphorylation as prerequisite to Thr172 AMPKα phosphorylation. In agreement, phospho-Ser65 PRKAG2 displayed a positive correlation with phospho-Thr172 AMPKα in PC patient tumors where both sites were downregulated compared to healthy human adrenal medullae. The interaction identified between PRKAG2 and AMPKα appears to be isoform-specific. Several mutations in PRAKG2 have been associated with Wolff-Parkinson-White syndrome, hypertrophic cardiomyopathy, and glycogen storage disease of the heart. Mutations such as K475E, and R302Q in *PRKAG2* gene lead to increased AMPK activity associated with cardiomyocyte hypertrophy (Xu et al., 2017; Zhan et al., 2018). However, the isoform-specific function of PRKAG2 has remained largely unexplored in cancer. PRKAG2 nuclear translocation in response to AMPK activation has also been shown to promote cardioprotection against ischemic injury (Cao et al., 2017). Nuclear-to-cytosol translocation of PRKAG2 and subsequent AMPK activation is likely due to increased interaction of the PRKAG2-AMPK complex with LKB1(Xie et al., 2008). Considering the varied functional aspects of PRKAG isoforms, the precise molecular mechanism by which GSK3-dependent Ser65 PRKAG2 phosphorylation mediates Thr172 AMPK phosphorylation and activation warrants further study and additional features of PRKAG2-AMPK regulation may also come to light.

AMPK-dependent signaling to p53 regulates the expression of multiple p53 targets, and subsequently affects cellular proliferation (Imamura et al., 2001). Here, Cdk5 inhibition activated AMPK, which in turn elevated Ser15 p53 phosphorylation and arrested cell cycle progression at G1 phase (Jones et al., 2005). Glucose deprivation or other forms of energetic stress may invoke AMPK-p53 signaling as a stress response mechanism. However, activation of AMPK-p53 signaling can also drive anti-glycolytic (*i.e.*, anti-Warburg) effects similar to that which we observed in response to Cdk5 inhibition (Thoreen and Sabatini, 2005). This is consistent with the ability of AMPK to negatively regulate glycolysis in cancer cells as a feature of its anti-tumor effects (Faubert et al., 2013). *SDHB* loss triggers a shift to high glycolytic phenotype, as seen in most PC patients (Her and Maher, 2015; Jochmanova and Pacak, 2016). Interestingly, inhibiting Cdk5 attenuates glycolysis, suggesting the kinase has an active role in maintaining aerobic glycolysis while maintaining low levels of AMPK-p53 signaling. *SDHB* deficiency can also manifest a hypermethylator phenotype by silencing tumor suppressor genes such as p16^INK4a^ (Kiss et al., 2013). Interestingly, here Cdk5 inhibition increased p16^INK4a^ expression, suggesting a possible role for Cdk5-AMPK signaling in regulation of demethylases as an epigenetic avenue that should be explored further (Rao et al., 2020).

In summary, this study reveals a novel phospho-dynamic mechanism where a Cdk5/GSK3/PRKAG2-AMPK/p53 signaling cascade acts as a critical downstream effector of SDHB loss. We demonstrated key components of this cascade across cell-based and *in vivo* models as well as in human tumors. We also derived a novel clinically accurate model of PC by transgenically invoking aberrant Cdk5 activity. These findings serve as a mechanistic rationale to utilize combinations of Cdk5 inhibitors and AMPK agonists in tumors driven by Cdk5/AMPK-dependent metabolic checkpoint.

## METHODS

### Cell culture, Reagents, Plasmids

#### Cells

The human progenitor PC cell line, hPheo1 (Ghayee et al., 2013) and mouse MTT PC cells were used (Korpershoek et al., 2012). SDHB was deleted from parent hPheo1 using CRISPR/Cas9 technology. (sgRNA sequence used was ATGGCAAATTTCTTGATACG and partial knockout clones were selected). Cells were cultured in RPMI 1640 with 10% FBS (Gibco). HEK293, HeLa, MDAMB-231, and H1184 were maintained in ATCC-recommended culture.

#### Pharmacologic inhibitors

Roscovitine, CVT313, MRT3-007, MRT3-124, CR8 were from Dr. Meijer (ManRos Therapeutics, France). 25-16 was from Dr. Natarajan (University of Nebraska), Metformin, and AICAR were from Sigma, NF-56-EJ40 (Med Chem Express# HY-130246).

#### Plasmids

Lentivirus gene expression vector (3rd generation; pLV-EGFP:T2A:Puro-EF1>ORF/FLAG) was used for subcloning ORF’s of PRKAG2 (NM_016203.3), DEPTOR(NM_022783.3), SDPR (NM_004657.5), AAK1(NM_014911.3) and UVRAG (NM_003369.3) driven by EF1A promoter. Cdk5 wild-type and kinase dead plasmids were described previously (Pozo et al., 2013).

### Database mining

SDHB copy number alteration, mRNA expression and co-expression analysis were analyzed in cBioportal Cancer Genomics Database (http://www.cbioportal.org). UALCAN online TCGA transcriptomic database was used to compare expression levels of SDHB/ Cdk5R1 between normal human medulla and PCPG patients (http://ualcan.path.uab.edu/). Publicly available Gene expression Omnibus (GEO) dataset GSE19422 was used for data mining in Figure S5. ShinyGO v0.61 gene-set enrichment tool was used to derive Gene Ontology categories, enriched pathways and graphical pathway network trees (Ge et al., 2020). Gene expression analysis of PRKAG2, AMPKα, (Figure S9D), Cdk5, Cdk2 and Cdk1(Figure S10A) were performed in GEPIA (The Gene Expression Profiling Interactive Analysis) database.

### Seahorse XF96 metabolic flux analysis

Real time extracellular acidification rate (ECAR) and oxygen consumption rate (OCR) in cells were determined using the Seahorse Extracellular Flux (XFe-96) analyzer (Seahorse Bioscience, MA, USA). 1×10^4^ cells were seeded per well of XF96 cell culture plates and incubated for 24 h to allow adherence. Bioenergetic profile was determined by mitochondrial and glycolytic stress tests following manufacture’s protocol. The following day cells were washed with pre-warmed XF assay base media (note: for OCR measurement, assay media was supplemented with 10 mM glucose, 1 mM Pyruvate, 2 mM L-glutamine, 5 mM HEPES and adjusted at 7.4 pH; for the glycolysis stress test, cells were washed with glucose-free XF base media). Cells were maintained in final volume of 180 μl/well of media at 37°C, in a non-CO_2_ incubator for 1 h. Meanwhile we loaded the cartridge ports with effectors. [For glycolysis stress test–12.5 mM Glucose, 1.5 µM Oligomycin, 50 mM 2- DG. For Mito stress test– 1.5 µM Oligomycin, 1 µM FCCP, 0.5 µM Antimycin and 0.5 µM Rotenone]. Measurements were normalized by protein content (BCA assay). Data were analyzed using XF96 Wave software and GraphPad Prism.

### Ca^2+^ measurement and live cell imaging

Cells were loaded with cell permeant intracellular Ca^2+^ flux indicator Fluo-4AM in HBSS (Gibco#14175) for 30 min following manufacturer’s instruction. Time-lapse live cell imaging was performed using Nikon A1R HD25 inverted confocal microscope equipped with perfect focus system. Cell imaging chambers were maintained in 5% CO_2_ and 37°C. Images were capture with Plan Apo λ 20× NA 0.8 wd 1000 objective, frame size– 1024×1024, scan speed– 2, time loop– 31, imaging zoom– 1.192, resolution– 3.7 pixels per micron, frame interval – 10 sec, bits per pixel–16, ex/em– 494/525. Pinhole and laser power settings were adjusted based on pre-stimulation background levels. Fiji software was used to select and quantify ROIs and individual object integrated density (IntDen) values were calculated for each cell using the formula (IntDen- (area of selected cell × mean field background intensity). Individual kinetic profile of [Ca^2+^]_i_ for WT and *SDHB* KO cells were generated by plotting fluorescence intensity as a function of time in seconds as described previously (Koh et al., 2016).

### Fluorescence resonance energy transfer (FRET)

CMV pCalpain-sensor (Addgene #36182) composed of eCFP (donor) and eYFP (acceptor) linked to calpain cleavage site (GSG-QQEVY GAMPRDGSG) where inactive calpains = High FRET and vice versa). pCalpain-sensor transfected in PC cells using Fugene HD according to the manufacturer’s instruction (Promega). Donor and acceptor bleed through were corrected using eCFP and eYFP only fluorophores. FRET measurements were performed using Nikon C2 confocal microscope (Plan Apo 60x Oil λS DIC N2) attached to a stage- top live-cell incubator to maintain cells at 37°C with 5% CO_2_ (Tokai Hit environmental chamber). FRET imaging based on sensitized emission were performed to acquire images with following filter combinations: donor (eCFP, ex-430±20, em-485±20); acceptor (eYFP, ex- 480, em-535±25); and FRET channel with a 435 nm (ex) and 535 nm (em). Visualization of FRET index and quantitation were performed as described in the FRET analyzer image J plug-in (Hachet-Haas et al., 2006).

### Tissue samples

Human PC were acquired from tumor bank of University of Alabama Birmingham, and National Institute of Child Health and Human Development (NICHD) following institutional review board (IRB) regulations, office for human research protections, NIH guidelines for research involving human subjects, and the health insurance portability and accountability act. All samples were de-identified, coded with no patient data, stored at -80°C until required for assay. Normal adrenal medullae were obtained from cadaveric kidney transplant donors (used as controls for which patients have given consent).

#### Immunostaining and Tissue microarray

Each tumor case was thoroughly reviewed, formalin-fixed, and paraffin-embedded blocks were acquired within Department of Surgery, University of Alabama at Birmingham, and Dept. of Pathology, NICHD followed by the protocol described previously using DAKO immunohistochemistry kit (Pozo et al., 2013). Human adrenal tumor tissue microarray (#AG801, US Biomax Inc.) was de-waxed at 60°C for 2 h followed by standard IHC protocol. Primary antibodies used included those for Cdk5 (Rockland# 200-301-163), -p35/25 (Cell Signaling#2680), ChrA (abcam#ab15160), anti-tyrosine hydroxylase (abcam#ab6211), and GFP (Cell signaling#2956). Secondary antibody alone was used as negative control. Quantitative analysis of DAB stained images were performed by using optical density (color deconvolution algorithm) within IHC profiler plug-in compatible with ImageJ digital image analysis software (Varghese et al., 2014).

### Pheochromocytoma animal model

All animal research was approved by the University of Alabama Birmingham (UAB) Institutional Animal Care and Use Committee (IACUC). Mice were genotyped and maintained within UAB Animal Resource Program Facility (ARP) and the animal research facilities of the Department of Surgery.

#### Transgenic model

PiggyBac technology was used to generate single-copy PNMT–tTA transgenic mice (C57BL/6) (Cyagen). Positive pups carrying PNMT-tTA transgene confirmed by genotyping using primers- Transgene PCR primer (Set1) Forward (F1): CAGTAGTAGATAAAGGGATGGGGAG; Reverse (R1): GGGGCAGAAGTGGGTATGATG; Annealing temp–60°C, product–350bp. Transgene PCR primer (Set2) Forward (F2)- CAGGAGCATCAAGTAGCAAAAGAG; Reverse (R2)- CACACCAGCCACCACCTTCT; Annealing temp– 60°C, Product– 373bp. All positive pups were confirmed by PCR to not contain any integration of the helper plasmid. Primers used in the PCR to test for helper plasmid integration, Forward–CTGGACGAGCAGAACGTGATCG; Reverse- CGAAGAAGGCGTAGATCTCGTCCTC. Bi-transgenic mice generated by crossing Tet-Op- p25GFP (Meyer et al., 2008) with that of PNMT-tTA. Transgenes were confirmed by genotyping for PNMT and p25-GFP alleles while control littermates were positive only for PNMT-tTA (Figure S6). Mice were treated with water containing 0.1 g/L doxycycline as required and all experimental mice were group-housed 12 h light/dark cycle with access to food and water *ad libitum*.

### Blood pressure

Mice blood pressure measurements were evaluated using CODA noninvasive BP system (a tail-cuff Method, Kent Scientific Corporation) as described previously (Wang et al., 2017).

### Assessment of Cdk5 inhibitors *in vivo*

#### Xenografts

6–7 week-old C.B-Igh-1b/IcrTac-Prkdcscid mice (Taconic Biosciences) were used for xenografts as previously described (Rai et al., 2020). 2 × 10^6^ hPheo1 tumor cells (WT and SDHB KO) were injected subcutaneously in the right flank of the mice. When average tumor volume reached 150 mm^3^, mice were divided into two groups and injected intraperitoneally with vehicle or MRT3-007 (0.5 mg/kg every alternate day for three weeks). Body weights and tumor diameters were measured 3 times/week and tumor volumes were calculated using the formula V = ab^2^ x 0.52, where a and b are major and minor axes of the tumor foci, respectively. The experiment was terminated on day 25, and the tumors were harvested for biochemistry and histological assessment.

#### Allograft and in vivo imaging

6-7 week old nude mice (nu/nu) (Jackson laboratory) were used for allograft assay described previously (Korpershoek et al., 2012). 1x10^6^ MTT luciferase expressing cells were injected via tail vein and imaged one-week post-injection via bioluminescence imaging (Xenogen IVIS). Cohorts bearing comparable size of allografts received IP injection of substrate d-Luciferin (250 μl;3.75 mg, Caliper Life Science, Hopkinton, MA, USA) 12 min before the whole-body imaging. Data was acquired and analyzed using the Live Imaging software version 3.0 (Caliper Life Science). Fourteen days after injecting tumor cells, all mice were divided into three groups, treated with Vehicle, MRT3-007 (0.75 mg/kg) or Roscovitine (50 mg/kg) every other day for 2 weeks. Animals were re-imaged to measure metastatic lesions via bioluminescence imaging.

### Site directed mutagenesis and transfections

DNA modifications were performed in lentiviral (pLV) expression vectors to generate phospho- and dephospho-mimetics for PRKAG2 (S-65D/E/A), DEPTOR (S-244D/E/A), SDPR(S- 288D/E/A), AAK1(S-676D/E/A) and UVRAG (T-518D/E/A) using Q5-site directed mutagenesis Kit (NEB). Manufacturer’s instructions for mutagenic primer design were followed, and mutations were confirmed by DNA sequencing. Transfections were carried out using FuGene-HD (Promega). Transfection solutions were prepared in Opti-MEM using 1:3 ratio of plasmid DNA to Fugene transfection reagent.

### Cell growth assay

Cells were seeded in 6-well plates at the density of 1x10^5^, transfected with phospho- or dephospho-mimetics and control vectors. Cell growth was determined 48 h post-transfection by dual fluorescence acridine orange/propidium iodide (AO/PI) viability staining. Total cell number was counted using Cellometer Auto 2000 cell viability counter (Nexcelom) and normalized with total number of GFP-expressing cells. Dose-response curves were generated by cell viability tests using Cell Counting Kit-8 (CCK-8) (Dojindo). Assays were performed in five replicates and repeated at least 3 times.

### Phosphorylation state specific antibody generation

Production and affinity purification of phosphorylation state specific polyclonal antibodies were performed as described previously (Hemmings, 1997). Phospho-Ser65 PRKAG2 were raised against a synthetic oligopeptide encompassing the amino acid sequence (RKVD<u>S*P</u>FGC). Cysteine containing phosphopeptide was conjugated to carrier protein *Limulus* hemocyanin using hetero-bifunctional crosslinker m-maleimidobenzoyl-N-hydroxysulfosuccinimide ester (Sigma# 803227). This conjugate was used to immunize New Zealand white rabbits (Charles River Laboratories). Preimmune sera were obtained and booster injections of 150 µg phosphopeptide conjugate were given at 2, 4, 6 and 8 weeks. Blood was collected at weeks– 5, 7, 9,11,13 and 14. The specificity of the antibodies in anti-serum was characterized by dot blot analysis using dephospho- and phospho-PRKAG2 standards. Phosphorylation-specific antibodies were purified using affinity purification method (Hemmings, 1997).

### Immunoblotting, Cell cycle analysis, and *in vitro* phosphorylation

Antibodies to the following phosphorylation sites and proteins were used: phospho-Thr172 AMPK (#2535), phospho-Ser79 ACC (#11818), phospho-Ser21/9 GSK3 (#9223), phospho-Ser16 p53 (#82530), p27Kip1(#3686), phospho-Ser139 H2Ax (#80312) from Cell Signaling Technology; anti-p16INK4A (Proteintech#10883-1-AP), anti-spectrin (Millipore Sigma#MAB1622) and, anti-GAPDH (#ZG003), anti-actin (#PA5-78715) from Thermofischer Scientific. SDS-PAGE and immunoblotting were conducted as previously described (Bibb et al., 1999). For cell cycle analysis, MRT3-007 treatment or phosphomimetics transfected cells were stained with 50 µg/ml of propidium iodide and analyzed for cell-cycle distribution as described previously (Erba et al., 1989). *In vitro* phosphorylation and immunoprecipitation-kinase assays were performed using optimized protocols described previously (Bibb et al., 1999; Pozo et al., 2013).

### Magnetic Resonance Imaging

MRI experiments were performed using a Bruker Biospec 9.4 Tesla scanner with Paravision 5.1 software (Bruker Biospin, Billerica, MA). A Bruker 72 mm volume coil was used for signal excitation, with a 24 mm surface coil for reception (Doty Scientific Inc., Columbia, SC). Mice were anesthetized with isoflurane gas and respiration observed with an MRI-compatible physiological monitoring system (SA Instruments Inc., Stony Brook, NY). Animals were imaged in supine position on an animal bed with integrated circulating heated water to maintain temperature during the experiment. Scout images were collected in the axial, sagittal, and coronal planes to confirm animal positioning and coil placement. A 2D T2-weighted fast spin echo sequence was used for imaging of kidney and adrenal gland areas. Prospective respiratory gating was enabled to minimize motion artifacts. The following imaging parameters were used: TR/TE=2000/25 ms, echo spacing=12.5 ms, ETL=4, 4 averages, 23 contiguous coronal slices with 0.5 mm thickness, FOV=30x30 mm, and matrix=300x300 for an in-plane resolution of 100 µ. All MRI images were obtained in the DICOM (Digital Imaging and Communications in Medicine) format and were imported into the image processing ITK software to obtain tumor volumes and perform 3D reconstructions. Mean tumor volumes were measured by drawing regions of interest (ROI), to circumscribe the entire tumor.

### Statistical analysis

Data were analyzed by one-way ANOVA combined with Tukey’s multiple-comparisons test or Student’s *t*-test using Prism 8 version 8.4.2 (Graph Pad Software). Statistical significance was defined as *p*-value *< 0.05, **p<0.01, ***p<0.001, ****p<0.0001.

## ACKNOWLEDGEMENTS

We thank Samuria Thomas for assistance with MRI, Daniel Epstein for technical expertise, Stephen Barnes for LC/MS succinate measurements, David Pollack for assistance in measuring mouse blood pressure, Haydn Ball and the University of Texas Southwestern Medical Center (UTSW) Protein Technology Core for peptide synthesis, Leslie Bibb and SouthernBiotech for generation of phosphorylation state-specific antibodies, and Pfizer for CP681301. This research was supported by the SDHB Para/Pheo Alliance and a Neuroendocrine Tumor Research Foundation (NETRF) Award (J.A.B.). Portions of this work were also facilitated by NIH support (MH116896, MH126948, J.A.B.), the Robert E Reed Gastrointestinal Oncology Research Foundation and an American Cancer Society Institutional Research Grant Junior Faculty Development Award (S.R.), and an American Cancer Society Postdoctoral Fellowship (A.M.C.). This research was supported, in part, by Intramural Research Program of the *Eunice Kennedy Shriver* NICHD, NIH (K.P.), the Gatorade Trust through funds by the University of Florida, Department of Medicine (H.K.G.) and by a EUROSTARS grant (CYST-ARREST, L.M.). Studies were also supported by the National Cancer Institute Cancer Center Support Grant P30 CA013148 (UAB O’Neal Comprehensive Cancer Center) and used the UAB High Resolution Imaging Facility and the UAB Preclinical Imaging Shared Facility. It was also supported by S10 OD028498-01 (UAB Preclinical Imaging Shared Facility). Some chemical synthesis was also supported by NCI SPORE P50 CA127297 (A.N., Eppley Institute for Research in Cancer and Allied Disease). This work was also supported in part by grant 1145265 from the National Health and Medical Research Council (NHMRC, to J.S.O.), St Vincent’s Institute of Medical Research (Australia) and the Victorian Government’s Operational Infrastructure Support Program. We are grateful to the UAB Diabetes Research Center (NIH P30 DK-079626) for providing outstanding core services in support of this research.

## SUPPLEMENTARY FIGURE LEGENDS

**Figure S1.**
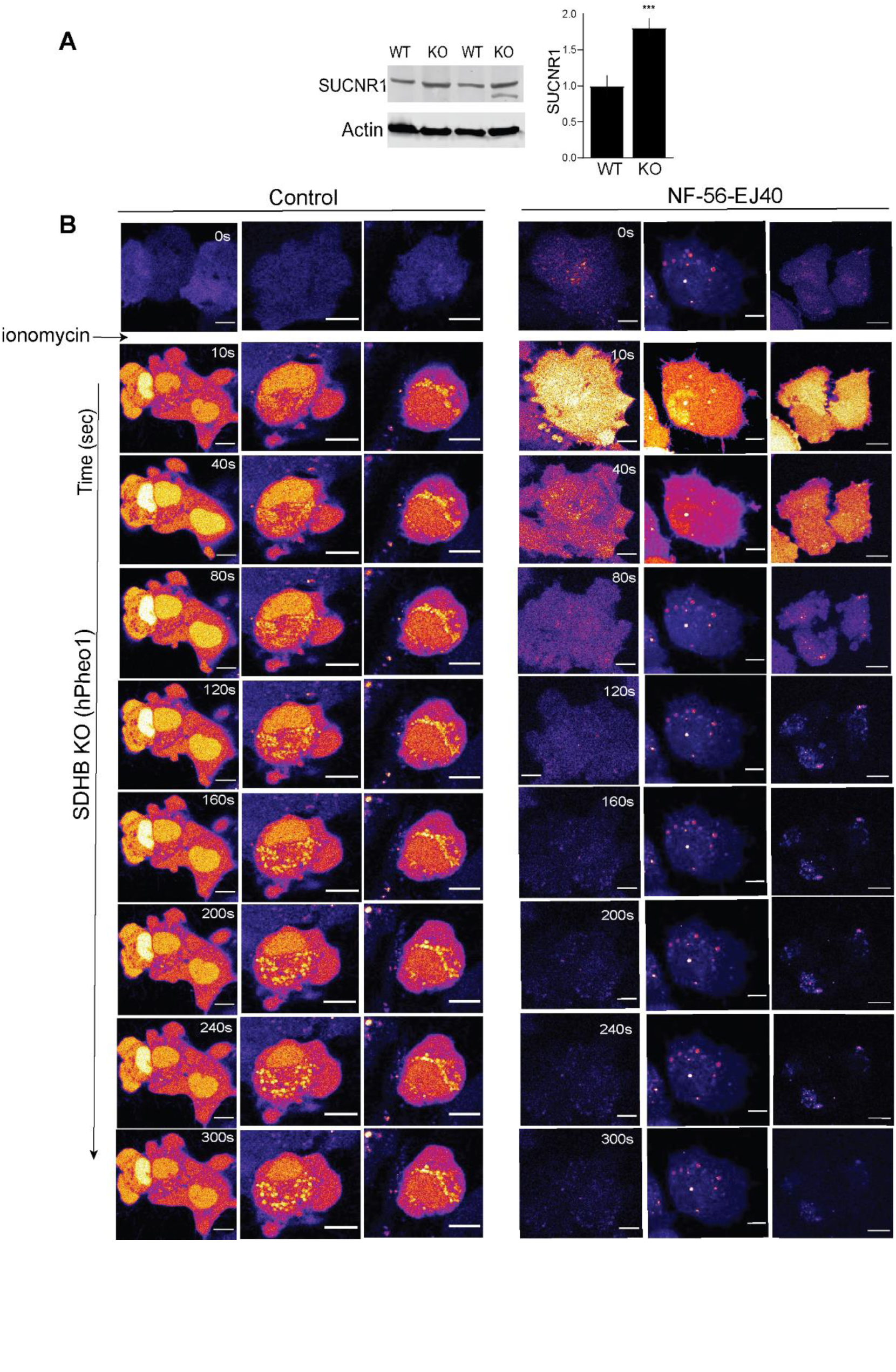
Loss of *SDHB* perturbs SUCNR1-mediated Ca^2+^ homeostasis (A) Relative SUCNR1 expression in WT vs*. SDHB* KO cells. Data are means ± S.E.M., ***p<0.001. (B) Supplemental confocal time lapse images showing [Ca^2+^]_i_ changes in control vs. NF-56-EJ40 treated cells as indicated in response to ionomycin (10 µM).

**Figure S2.**
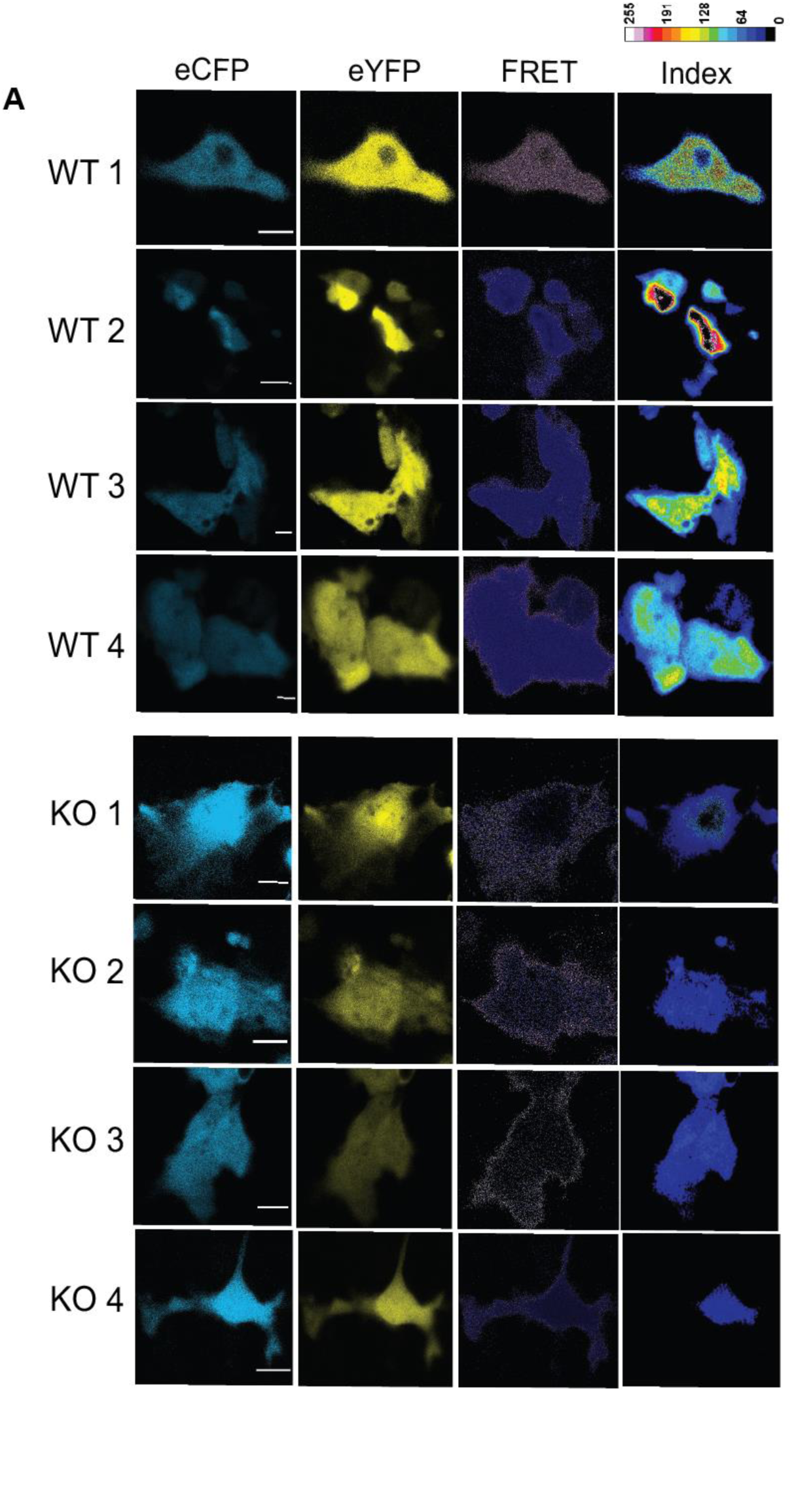
FRET sensor-based visualization of calpain activity in PC cells. (A) Representative live cell confocal photomicrographs shows donor (eCFP), acceptor (eYFP) and ratiometric FRET index. High FRET efficiency is detected in WT (images 1-4) compared to KO PC cells (KO 1-4).

**Figure S3.**
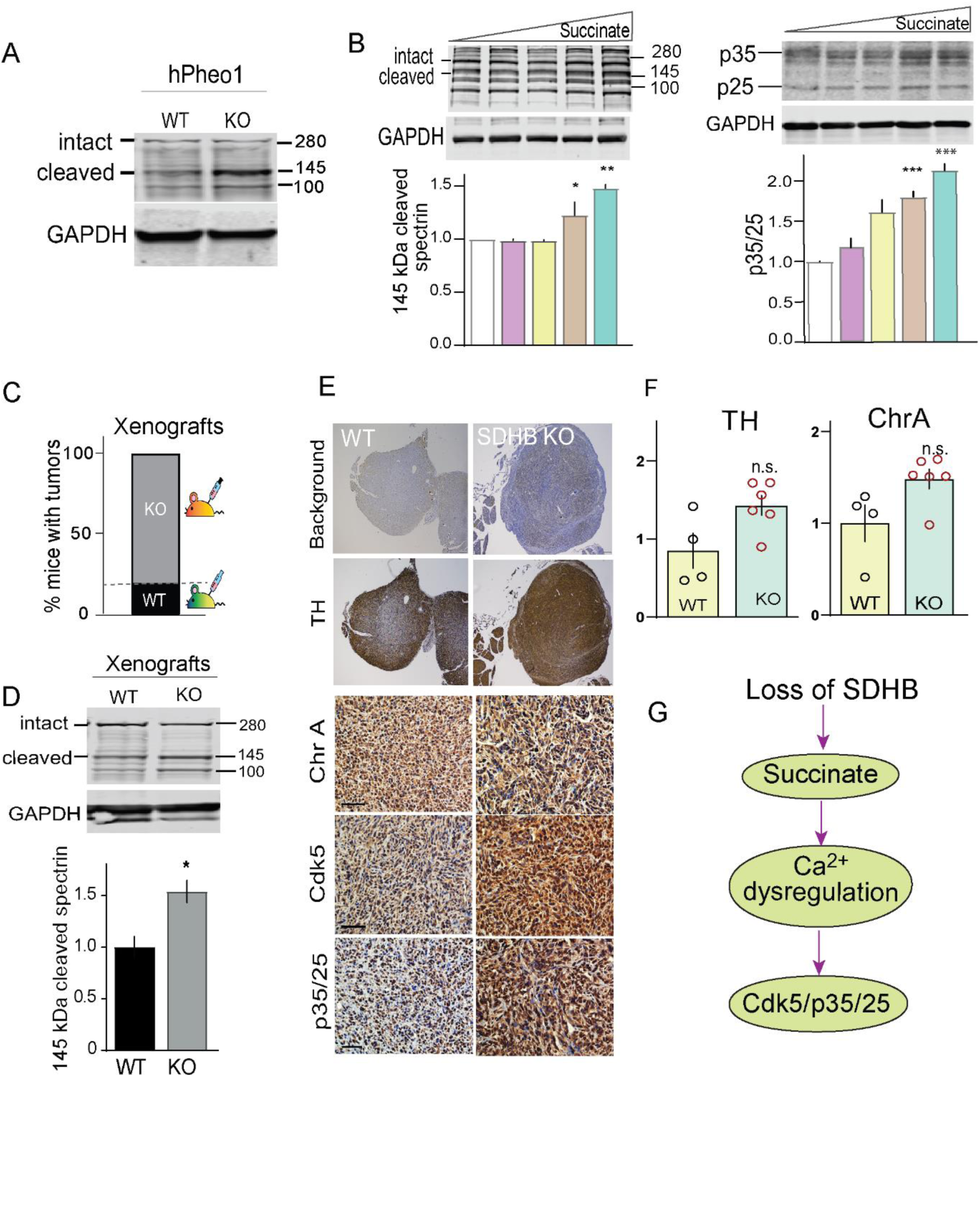
Characterization of *in vitro* and *in vivo* model of hPheo1. (A) Immunoblot showing spectrin cleavage as a function of calpain activity in PC cells. (B) Quantitative immunoblots showing dose-dependent effect of dimethyl succinate (0, 0.5, 2.5, 5, 10 mM) on spectrin cleavage and p35/25 levels in WT hPheo1. (C) Plot comparing % tumor burden of WT vs. KO hPheo1 derived xenografts. (D) Quantitative immunoblotting of spectrin cleavage in extracts derived from WT (n=4) and KO (n=6) xenografts. (E) Immunostains for chromaffin cell neuroendocrine markers, TH, ChrA, Cdk5 and p35/25 in xenograft tissues. (F) Immunoblot quantification of TH, and ChrA in WT and KO tumor tissues as indicated. (G) Signaling schematic showing aberrant Cdk5 as a downstream effector in response to SDHB loss. n = 11 (C) and 4-6 (F), data are means ± S.E.M., *p < 0.05, **p < 0.01, ***p<0.001, n.s. non-significant, Student’s *t*-test or one way ANOVA.

**Figure S4.**
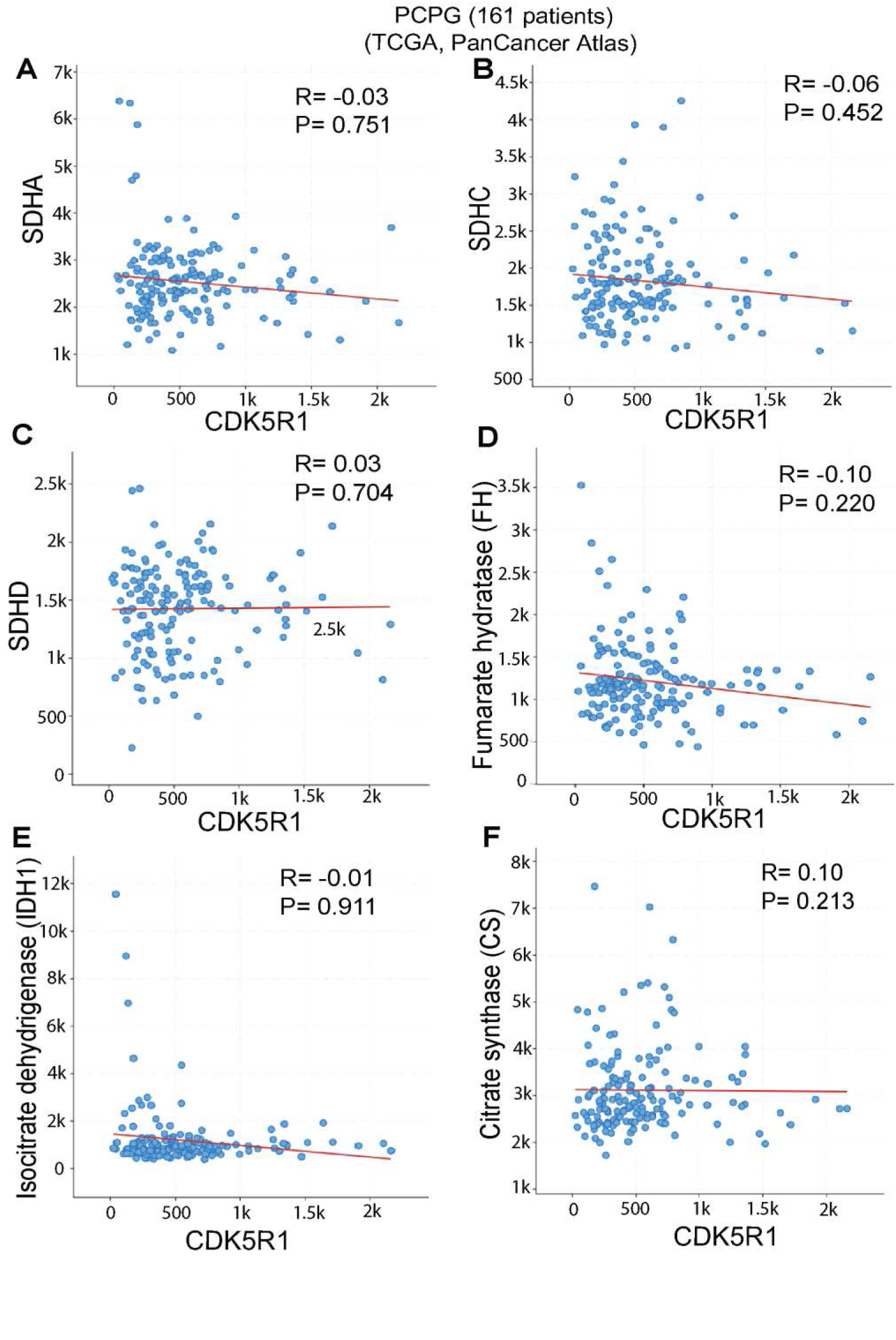
Gene expression correlation analysis for tumor suppressive TCA genes with CDK5R1(p35) using cBioPortal. Scatter plot shows Spearman’s correlation of CDK5R1 with (A) SDHA, (B) SDHC, (C) SDHD, (D) FH (Fumarate hydratase), (E) IDH1 (isocitrate hydratase), (F) Citrate synthase (CS).

**Figure S5.**
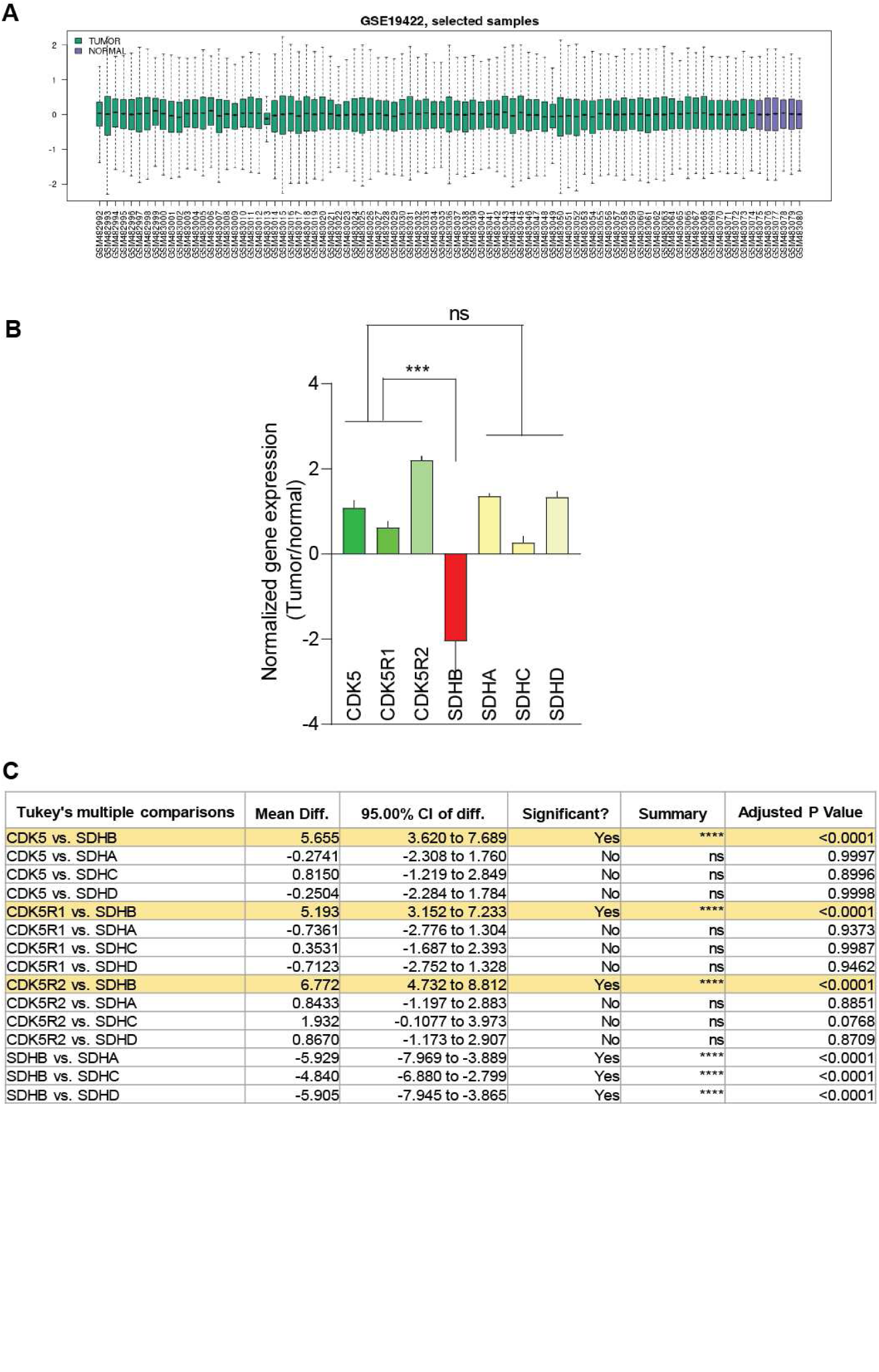
Gene expression analysis of Cdk5/SDHB signaling components in human PCPG dataset. (A) Box plot showing distribution of gene expression values for PC (n=84) and normal adrenal tissues (n=5) following normalization (NCBI/GEO/GSE-19422). X and Y-axis represents GEO samples and expression values. (B) Plots comparing gene expression values of Cdk5 signaling components with that of SDHB, SDHA, SDHC, and SDHD. (C) Table presents the statistical significance following analysis of gene expression values comparing Cdk5 signaling components with that of SDHB, SDHA, SDHC, or SDHD as indicated. Data are means ± SD ***p<0.001, n.s. non-significant, Tukey’s multiple comparisons test.

**Figure S6.**
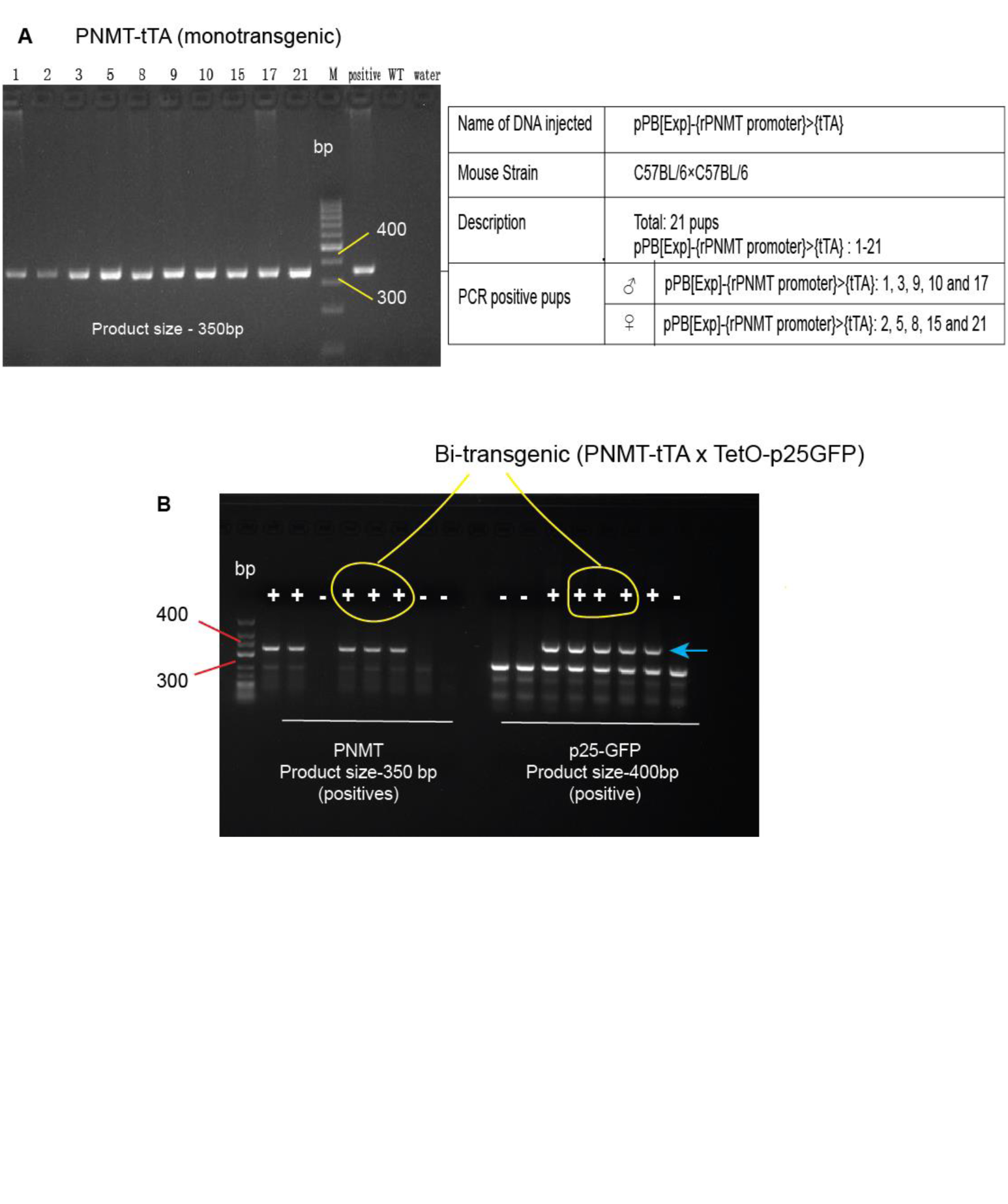
Genotyping of bitransgenic mice. (A) PNMT–tTA littermates were screened to identify positive progeny following PCR analysis using primers described in methods. Out of 21 pups screened, 10 were positive for PNMT-tTA gene as shown in the table (right). (B) Bi-transgenic pool (PNMT-tTA×TetO-p25GFP) were identified by detecting transgenes expression of PNMT (350 bp) and p25-GFP (400 bp) by RT-PCR.

**Figure S7.**
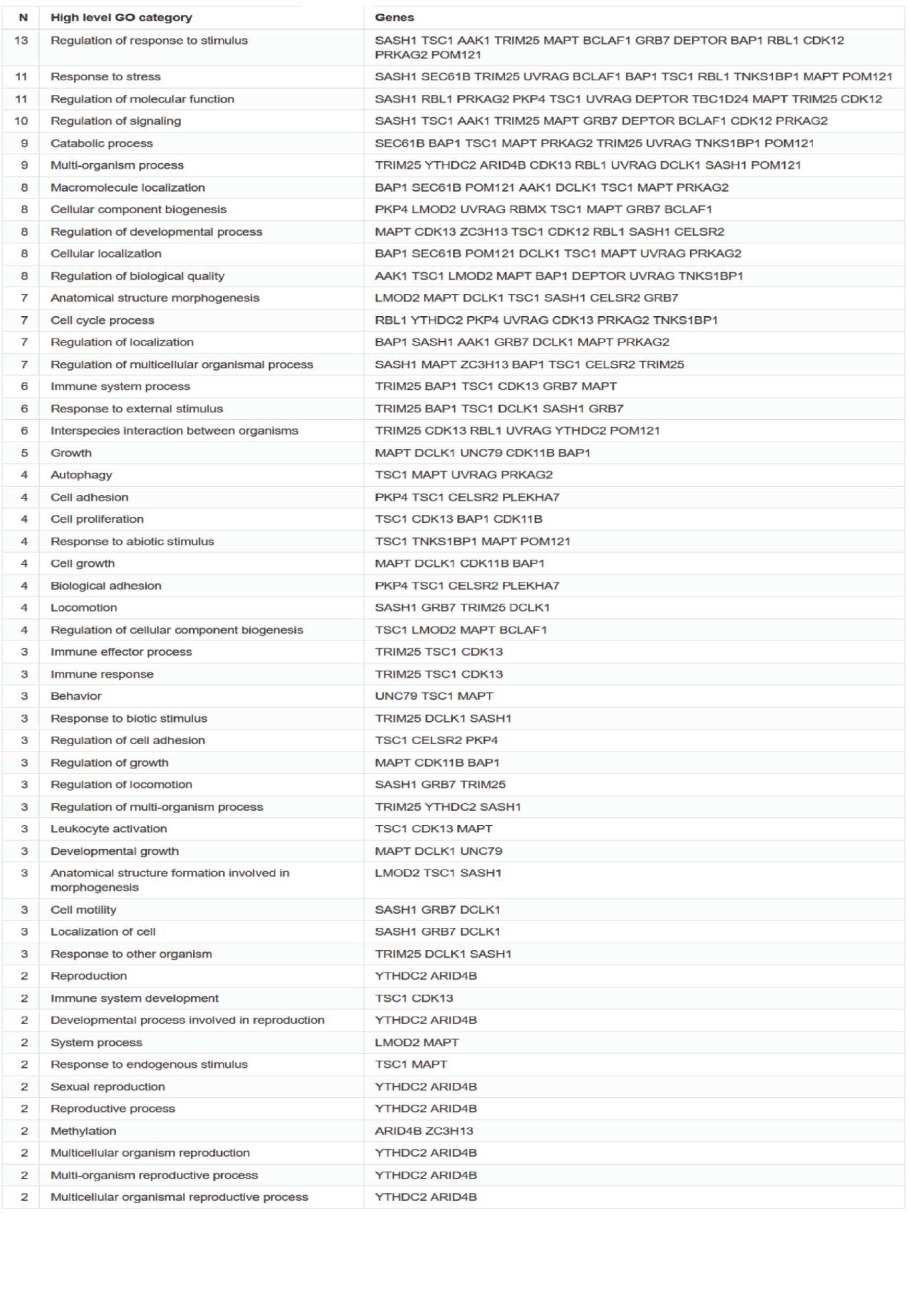
Table showing enriched GO category (biological processes) corresponding to the downregulated Cdk5 targets in p25OE tumors (data derived from ShinyGO web tool, followed by FDR correction; p-value cutoff = 0.05).

**Figure S8.**
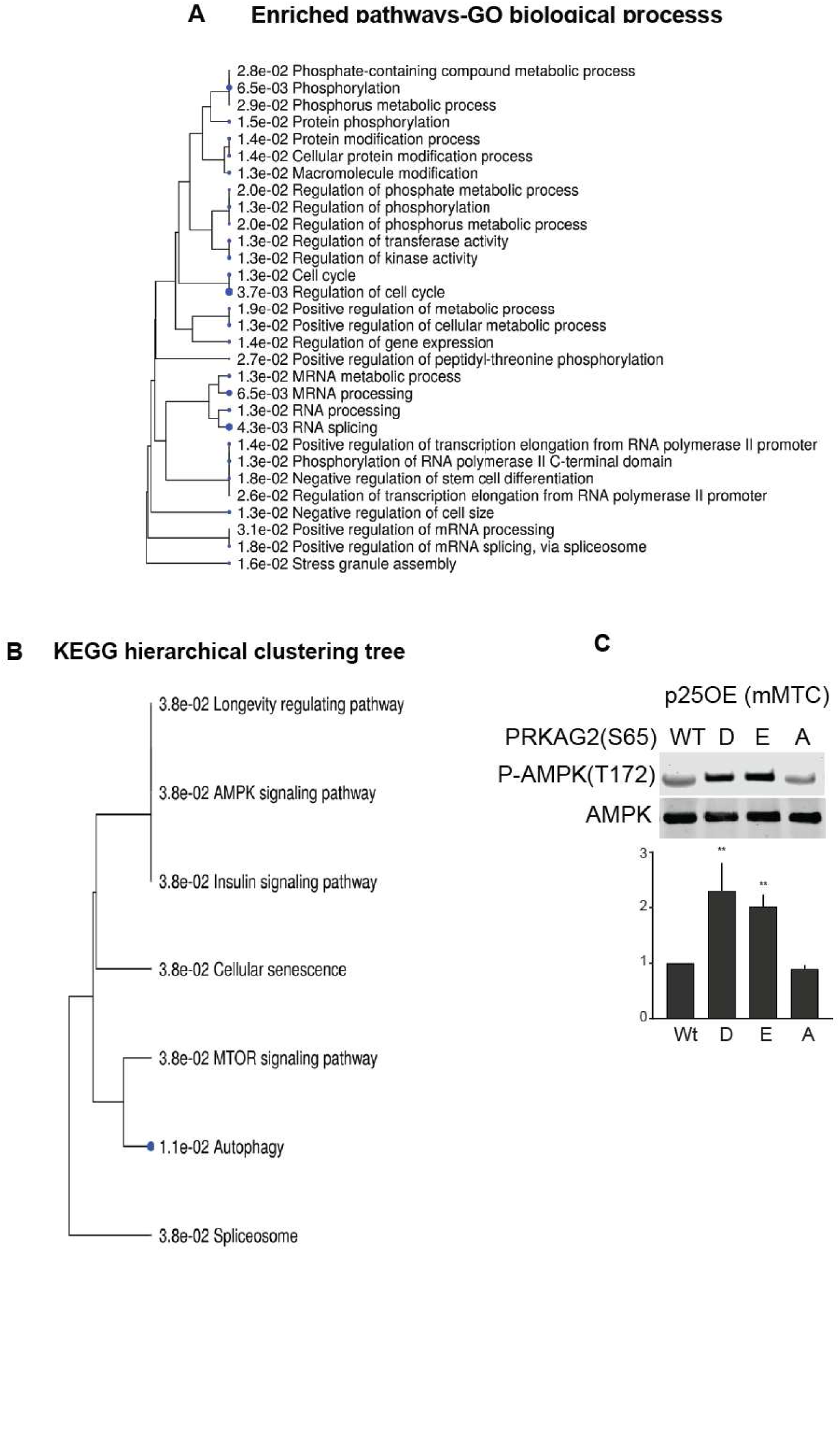
Relation among enriched GO molecular pathways visualized as (A) Hierarchical clustering tree, and (B) Kyoto Encyclopedia of Genes and Genomes (KEGG) pathway analysis. Downregulated phospho-target genes mostly involved in cellular metabolic and protein phosphorylation processes as shown. (C) Quantitative immunoblotting of (P-T172) AMPK catalytic phosphorylation in response to PRKAG2 S65D/E/A phosphosite mutant expression in p25OE mMTC cells (mouse <u>M</u>edullary <u>T</u>hyroid <u>C</u>arcinoma cells). Data are means ± SEM, **p<0.01, n=3.

**Figure S9.**
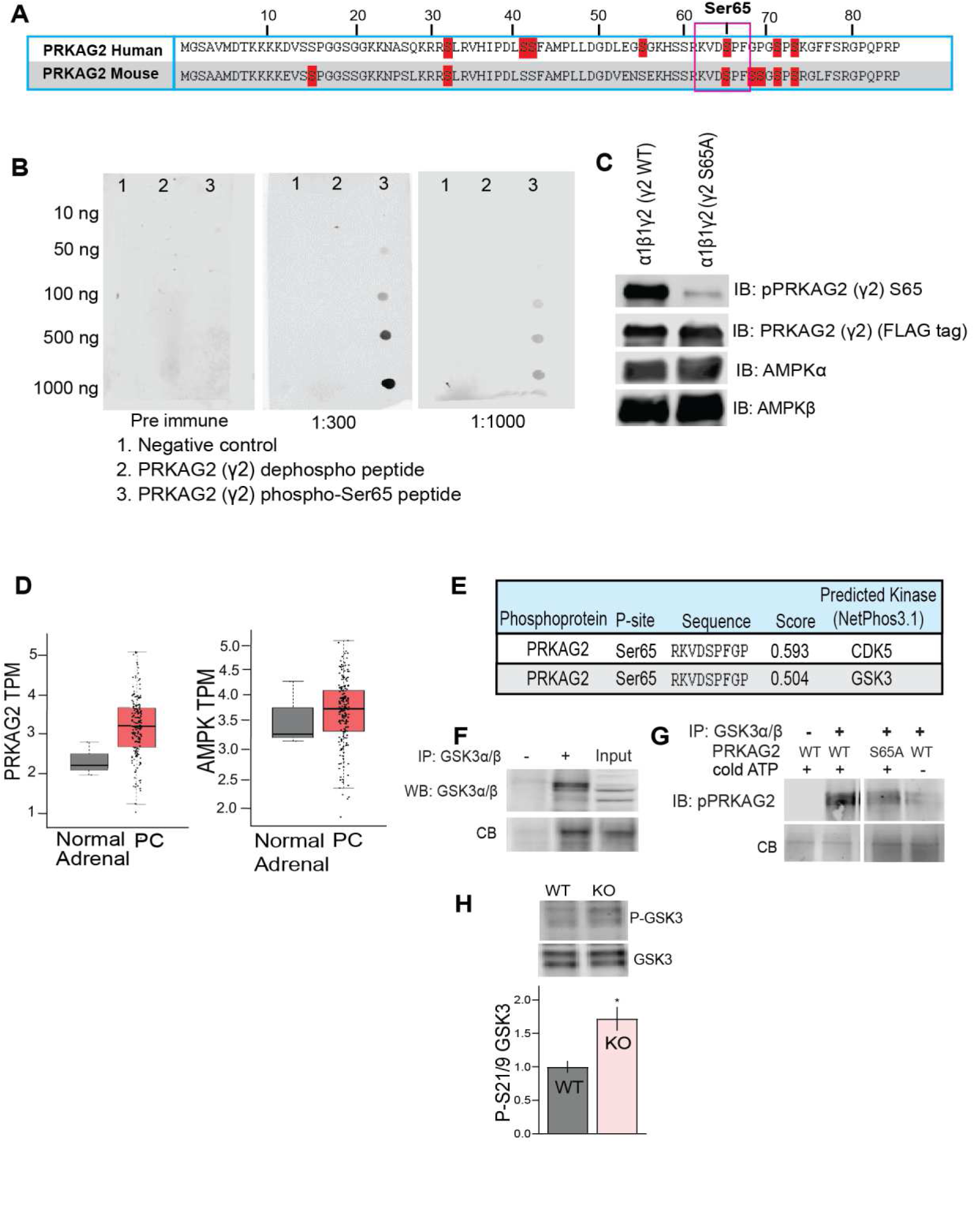
Generation of phospho-Ser65 PRKAG2 antibody. (A) Clustal omega alignment shows conservation of Ser65 phosphorylation site in human and mouse PRKAG2 amino acid sequence. (A) Dot blots shows selective reactivity of anti-serum for phospho-PRKAG2 peptide. (C) Immunoblots of AMPK holo-complex (α1β1γ2) comprised of WT or S65A PRKAG2 phosphorylated by GSK3β *in vitro,* demonstrating specificity of affinity purified antibody for P-S65 PRKAG2 (WT) and not the S65A form. (D) The mRNA expression of PRKAG2 and AMPKα in PC vs. normal adrenals. Data derived from Gene Expression Profiling Interactive Analysis (GEPIA) PCPG dataset, p-value cutoff 0.05. (E) NetPhos3.1 analysis suggests CDK5 and GSK3β as possible kinases that phosphorylates Ser65 PRKAG2. (F-G) IP kinase assay performed to phosphorylate purified PRKAG2 (WT, S65D, S65A) incubated with immunoprecipitated GSK3 (left) in the presence of cold ATP at 30°C for 60 min. The reaction products were probed with antibody to phospho-Ser65 PRKAG2 (right), CB=Coomassie blue stained protein. (H). Immunoblots densitometry shows differential expression levels of P-GSK3 in WT vs*. SDHB* KO cells.

**Figure S10.**
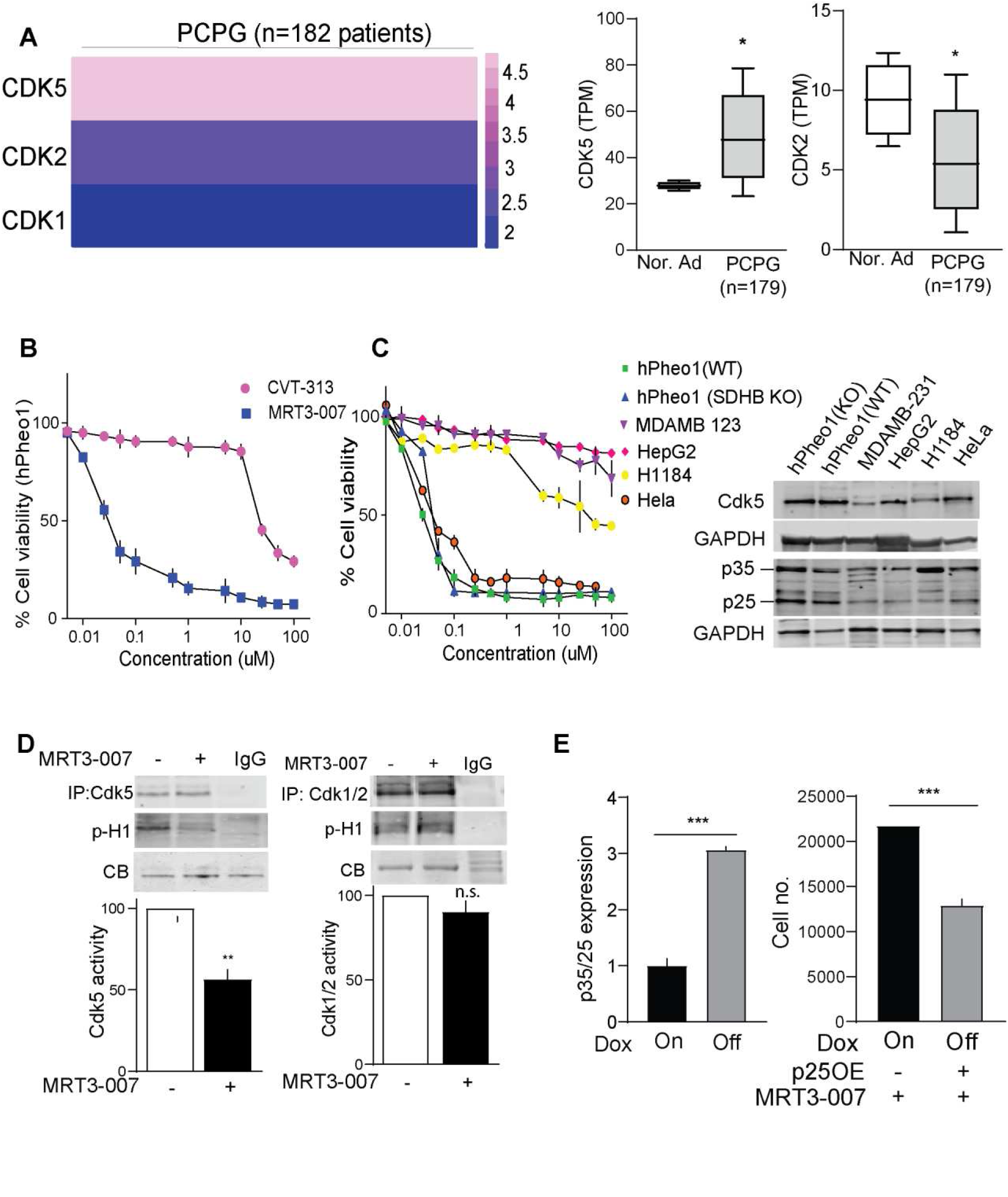
Selectivity assessment of MRT3-007 (Cdk5_in_). (A) Heatmap shows gene expression of Cdk5, Cdk2 and Cdk1 in human PCPG dataset analyzed in GEPIA (left). Bar graphs compares Cdk5/Cdk2 transcript levels in normal adrenals vs. PC tumors (right). (B) Growth inhibitory effects of Cdk5_in_ (MRT3-007) compared to that of the Cdk2 inhibitor (CVT-313) on hPheo1. (C) Dose-dependent effect of MRT3-007 on different cancer cell types including WT and *SDHB* KO hPheo1, MDAMB-231, HepG2, H1184, and HeLa (left) expressing high and low levels of Cdk5/p35/25 as indicated by immunoblot in right. Note MRT3-007 was least effective in cells expressing lowest levels of p25 (MDAMB-231, HepG2, and H184). Data are means ± S.E.M., n=3. (D) IP kinase assays with immunoblot for histone H1 phosphorylation (p-H1) with Cdk5 (left) and Cdk2 (right) immunoprecipitated from hPheo1 cell lysates treated with vehicle (-) vs. MRT3-007 (+) for 12 h. Data are means ± S.E.M., **p<0.01, n.s. non-significant, Student’s *t*-test, n=2. Coomassie blue (CB) staining shows total histone H1 protein levels. (E) Immunoblot quantitation confirming elevation of p25 in Dox OFF cells (left) as shown with effect of MRT3-007 (Cdk5_in_) on cell viability of p25-ON (Dox OFF) vs. p25-OFF (Dox ON) NE cells (right). Data are means ± S.E.M., ***p<0.001, Student’s *t*-test, n=3.

**Figure S11.**
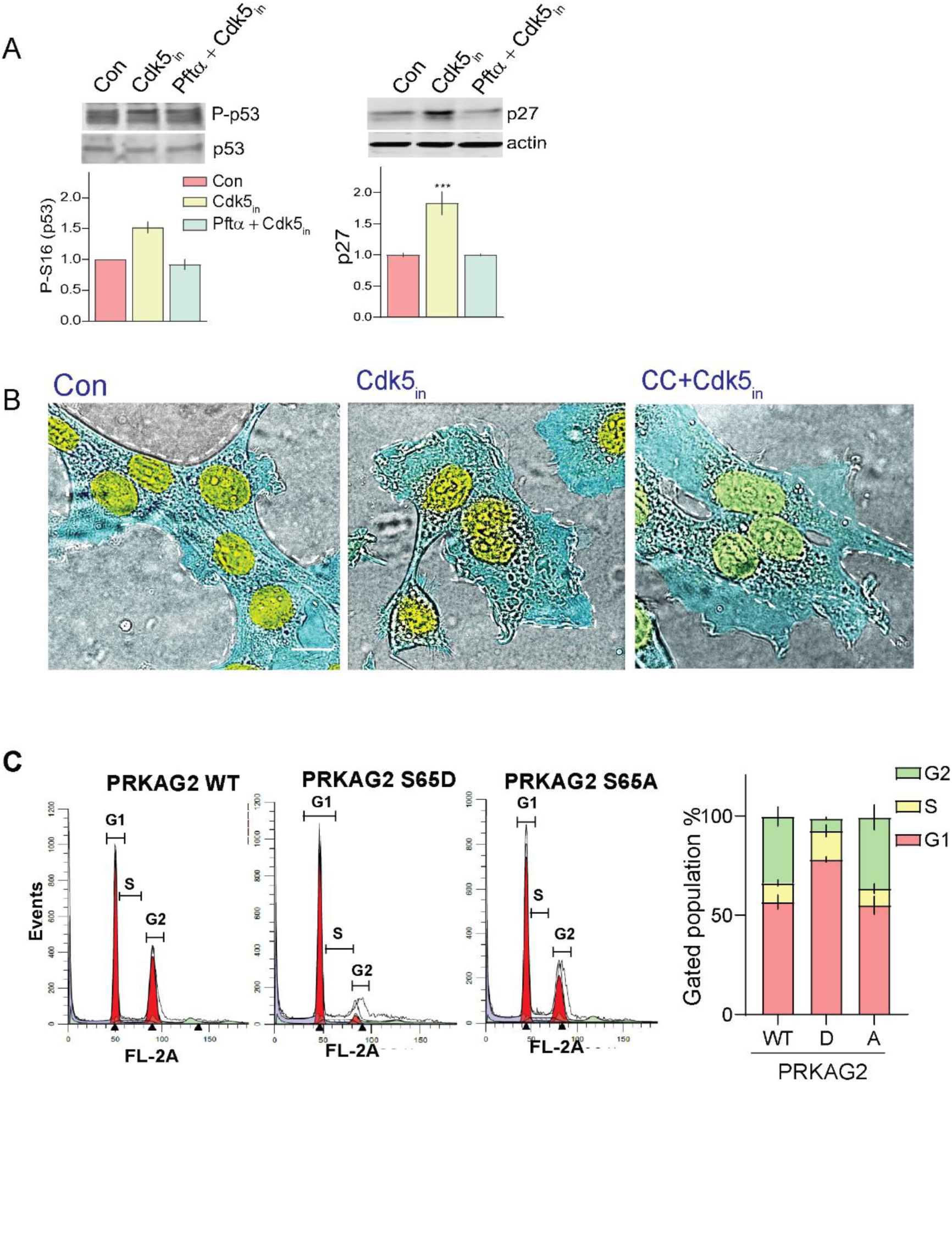
Inhibition of p53 or AMPK abrogate effects of Cdk5_in_ (MRT3-007). (A) Representative blots and densitometry detects P-p53 (S16) and p27^Kip^ upon Cdk5_in_ treatment in presence or absence of P53 inhibitor PFT-α (10 µM, 10 h pre-treatment). (B) Representative phase contrast microphotographs showing effects of AMPK inhibitor, Compound C (CC, 10 µM) on Cdk5_in_ induced morphological changes. (C) Representative histograms and quantitation indicating cell cycle profile of KO cells expressing either WT, S65D or S65A phosphomutants of PRKAG2.

**Figure S12.**
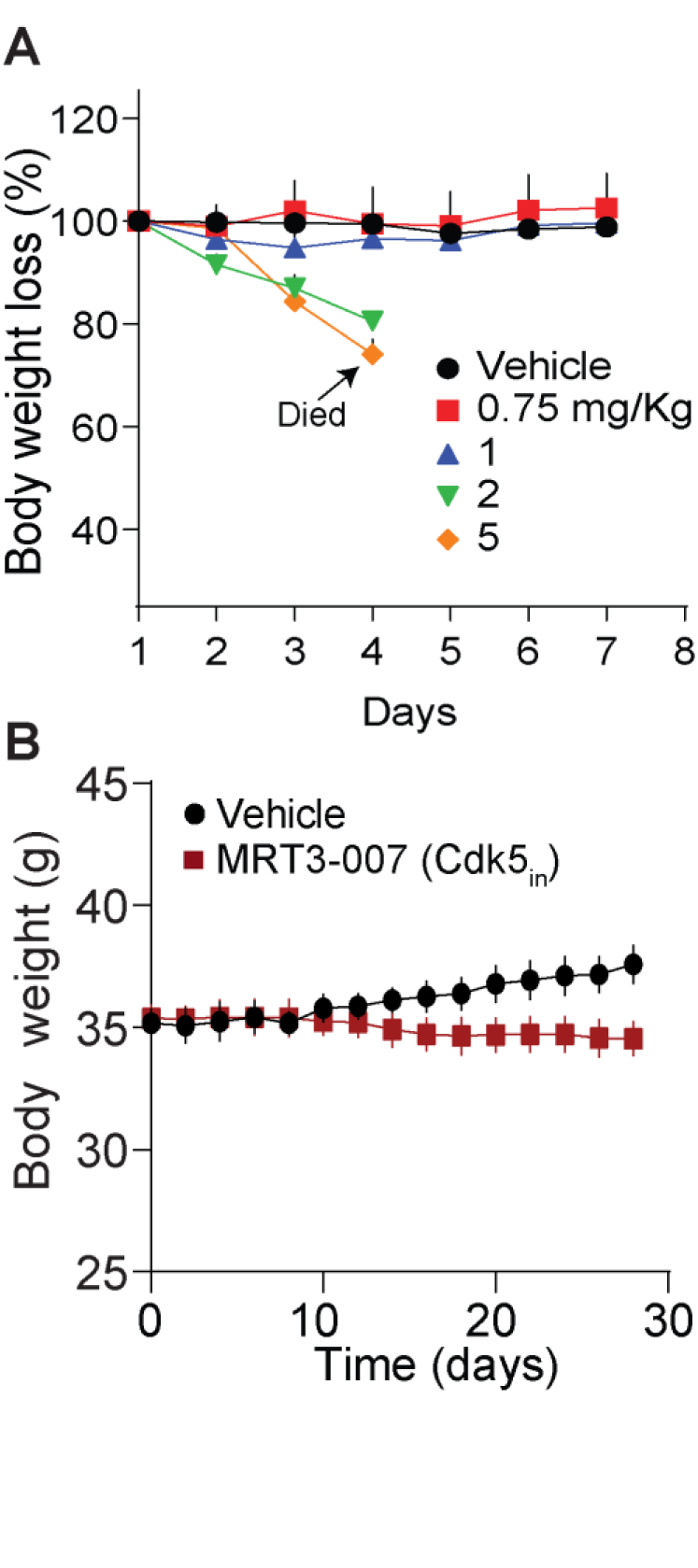
(A) Maximum tolerated dose (MTD) of Cdk5_in_ MRT3-007 in C57BL6 mice expressed as body weight (% of original). Mice injected with escalating doses of 0.75, 1, 2, or 5 mg/kg MRT3- 007, respectively (n=5). (B) Time-dependent effect of vehicle or MRT3-007 (0.5 mg/kg) on body weight of SDHB KO xenografts (n=4).

## Notes

### Competing Interest Statement

The authors have declared no competing interest.

